# Perturbation Curve models continuous transcriptional response trajectories and improves prediction of genetic modulations

**DOI:** 10.64898/2026.06.16.732192

**Authors:** Yunhua Zhong, Lifei Wang, Gui Yang, Lequan Yu, Xiaojuan Qi, Haibo Jiang

**Author notes:** These authors contributed equally: Yunhua Zhong, Lifei Wang. Correspondence should be addressed to Haibo Jiang, Xiaojuan Qi.

## Abstract

Single-cell CRISPR screens, Perturb-seq, have revolutionized functional genomics by revealing biological causality. However, although perturbation assignments are typically represented as discrete labels, the cell-level effective strength of perturbations is often continuous and diverse. Current analytical frameworks struggle to decouple the variability in perturbation strength from the diversity of downstream responses. Here, we present Perturbation Curve (PertCurve), a nonlinear, curve-based computational framework that models the trajectories of transcriptomic responses by explicitly incorporating diverse perturbation magnitudes and strengths. By ordering cells by perturbation strength, we demonstrate that PertCurve accurately recapitulates the response magnitudes and reveals the distinct modularity and asynchrony patterns of downstream gene behaviors. These patterns are categorized into archetypes, including proportional, sensitive, and threshold responses. By applying this framework across CRISPRi/a modalities, we identify universal response patterns in viral infection, apoptosis, and proliferation genes, and reveal previously overlooked context-specific regulatory features in cell differentiation. Finally, incorporating PertCurve into perturbation prediction models and evaluation metrics enhances predictive performance, delivering actionable insights for refining established models.

## Introduction

The advancement of various omics approaches, particularly single-cell omics methods such as single-cell transcriptomics and epigenomics, has generated vast amounts of data, substantially expanding our understanding of biological systems. However, the majority of these omics data remain descriptive, whereas deciphering biological systems fundamentally requires causal evidence. Perturbation experiments, which involve genetic or chemical interventions followed by measurement of cellular responses^1,2^, offer a powerful means to uncover causal relationships^3^. Perturb-seq integrates genome editing with single-cell sequencing: by precisely modulating target genes using CRISPR technology^4–8^ and subsequently profiling the transcriptional response of individual cells via single-cell RNA sequencing, it is especially well-suited for dissecting complex biological systems^9,10^.

One critical step in Perturb-seq analysis is to quantify two distinct layers of heterogeneity inherent to the datasets^4,5^. These layers encompass heterogeneity at the population level where different perturbed genes display varying overall perturbation strengths^4,5,11,12^, and at the single□cell level where cells subjected to the same perturbation exhibit diverse responses due to biological variability^4,5,13,14^. Furthermore, cellular-level heterogeneity encodes information about how gradual shifts in perturbation strength remodel downstream transcriptomic states^13,14^. The quantification of cellular-level and population-level heterogeneity represents two complementary analytical perspectives on the same dataset. An ideal analytical framework would integrate both aspects holistically. However, existing methodologies often address these aspects separately: some are designed to quantify heterogeneity in overall response strength across different gene perturbations^11,12^, while others focus on dissecting variable perturbation strengths at the single-cell level within the same targeted gene^13,14^. In addition, many response genes exhibit nonlinear dose–response relationships with gradual changes in perturbation strength^15–17^. Current methods that infer such relationships frequently rely on linear assumptions, which are fundamentally misaligned with the underlying nonlinear biology^13,14^. Dissecting this continuous perturbation strength and transcriptional response relationship is particularly valuable for studying complex disease-associated non-coding genetic variants, which typically act through moderate, allele-specific expression changes rather than complete deficiency effects.

Here, we developed Perturbation Curve (PertCurve), a nonlinear curve fitting method tailored for Perturb-seq data analysis. We demonstrate that PertCurve effectively outperforms conventional Perturb-seq data analysis methods across all respective analytical levels. By integrating cross-scale heterogeneity, PertCurve redefines normalized perturbation strength, enabling characterization of the biological significance of gene perturbations that prioritize transcription-regulating genes. Through the integration of CRISPRi and CRISPRa perturbation analysis with PertCurve, our method reveals characteristic universal and context-specific downstream transcriptomic response trajectory patterns as a function of perturbation strength, along with their underlying biological implications. Finally, incorporating PertCurve into state-of-the-art perturbation prediction models and evaluation metrics enhances predictive performance, delivering actionable insights to refine established models and develop new approaches.

## Results

### Overall Workflow

Perturb-seq with CRISPRi and CRISPRa exhibit heterogeneity at the population level and the single□cell level **(Fig. 1a**). To model this, we treat the collection of cells from single-gene perturbations and controls as a point cloud in a high-dimensional space. We posit that the distribution of these points arises from the superposition of the specific genetic perturbation and other sources of variability. The effect of perturbation can be represented as a one-dimensional manifold that admits fitting via a smooth curve embedded within the ambient high-dimensional space. We leverage prior knowledge of control/perturbation grouping information to anchor the curve direction and accurately fit the continuous dose-response curve from the baseline to the maximum perturbation effect. After fitting, each cell is projected onto the curve, effectively denoising the data by aligning cells along the estimated response path **(Fig. 1a and 1b)**. We term this approach Perturbation Curve (PertCurve).

**Figure 1.**
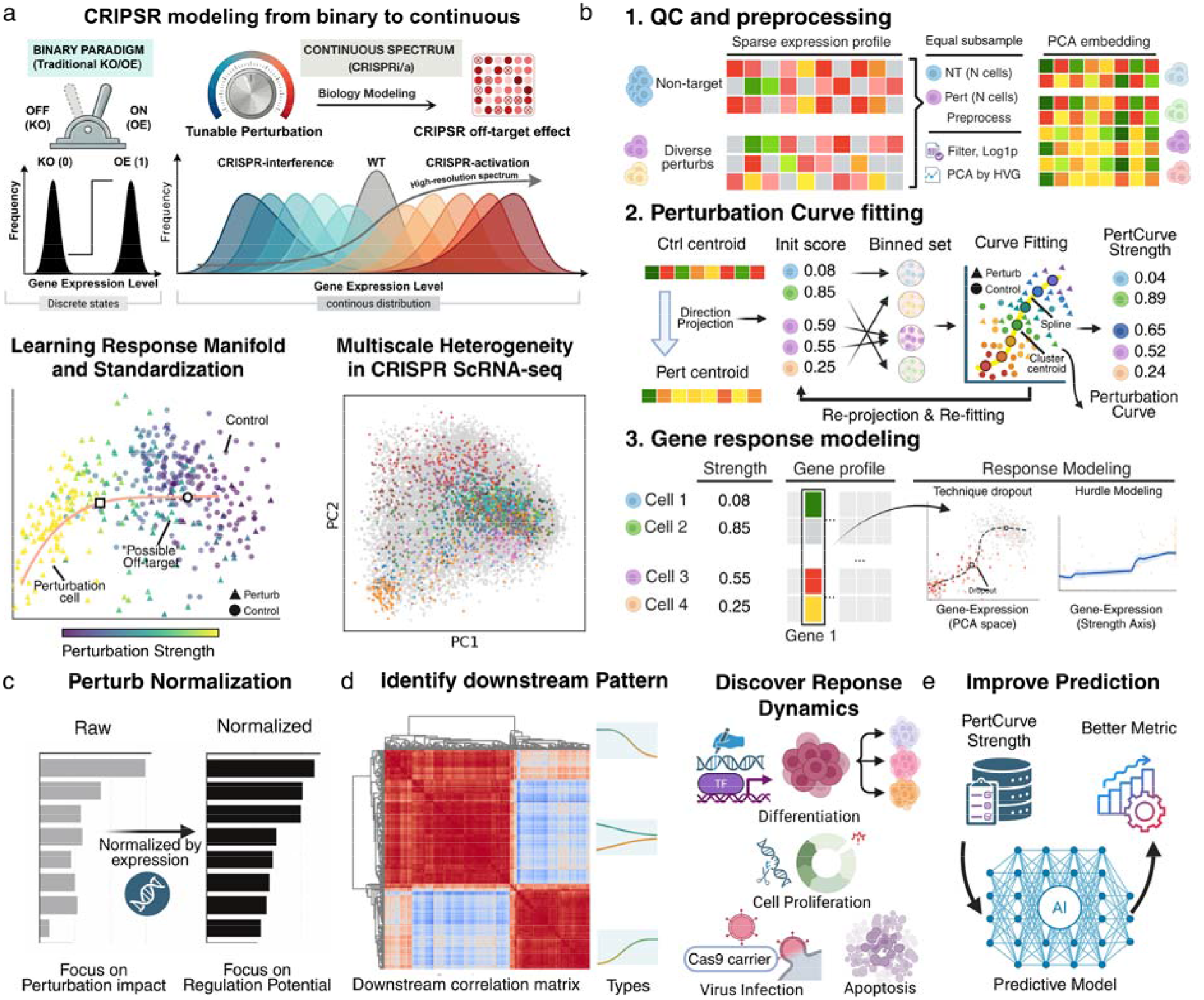
Overview of the Perturbation Curve (PertCurve) workflow. (a) Multilevel diversity in CRISPR screens. CRISPR genetic modulations exhibit heterogeneity at both the population and single-cell levels. *Top*, schematic of sources of diversity. *Bottom left*, PertCurve fitting for one perturbation, with colors indicating cell-wise perturbation strength. *Bottom right*, PCA of single-cell perturbation data showing diversity among targeted cells and different perturbation. Every color represent a perturbation population. (b) Schematic of the PertCurve workflow, including data preprocessing, curve fitting and response modeling. (c–e) Downstream applications of PertCurve in calculating normalized perturbation strength using population- and single-cell level heterogeneity derived from PertCurve (c); uncovering both universal and context-specific patterns of downstream gene responses across CRISPRi and CRISPRa perturbations (d); and enhancing perturbation-response predictive modeling (e).

Based on the PertCurve framework, we can quantitatively measure two levels of heterogeneity: (1) population-level heterogeneity across different perturbations, and (2) individual-cell-level transcriptomic response heterogeneity within the same perturbation condition **(Fig. 1b)**. These quantitative metrics enable a more refined quantitative analysis of perturbation on its target gene and the downstream response trajectories. For perturbation target genes, PertCurve normalized perturbation strength to measure their impact on the entire transcriptome **(Fig. 1c)**. For downstream response genes, PertCurve dissects the continuous perturbation strength and transcriptomic response relationship patterns, demonstrating certain genes respond universally to CRISPR genetic modulation experiments and revealing context-specific response trajectories along with their associated biological significance **(Fig. 1d)**. In addition, PertCurve can be incorporated for the development of perturbation prediction models, laying a solid foundation for both building new models and improving existing ones **(Fig. 1e)**.

### PertCurve quantifies multi-level heterogeneity in genetic perturbations and normalizes the perturbation strength of perturbed genes

For both CRISPRi and CRISPRa, a curve can be independently fitted to characterize the transcriptional response of cells following perturbation of a given gene (**Fig. 2a and 2b**). After fitting a curve to the transcriptional response of cells subjected to perturbation of a specific gene, we employed Wasserstein distance between perturbed cell population and control cell population on the curve to assess the overall strength of the perturbation response (**Fig. 2c**). For comparison, we benchmarked these metrics against the E-distance metric proposed in the scPerturb framework^12^ across both the Replogle (CRISPRi) and Norman (CRISPRa) datasets (**Supplementary Fig. 1**). Because our metric is inherently normalized, it is robust to variations in sample size and sequencing depth, enabling direct cross-modality comparisons and integrated analyses across perturbation modalities. In contrast, the E-distance metric scales with technical factors, resulting in pronounced discrepancies across different experimental platforms even under identical preprocessing pipelines (**Fig. 2d**).

**Figure 2.**
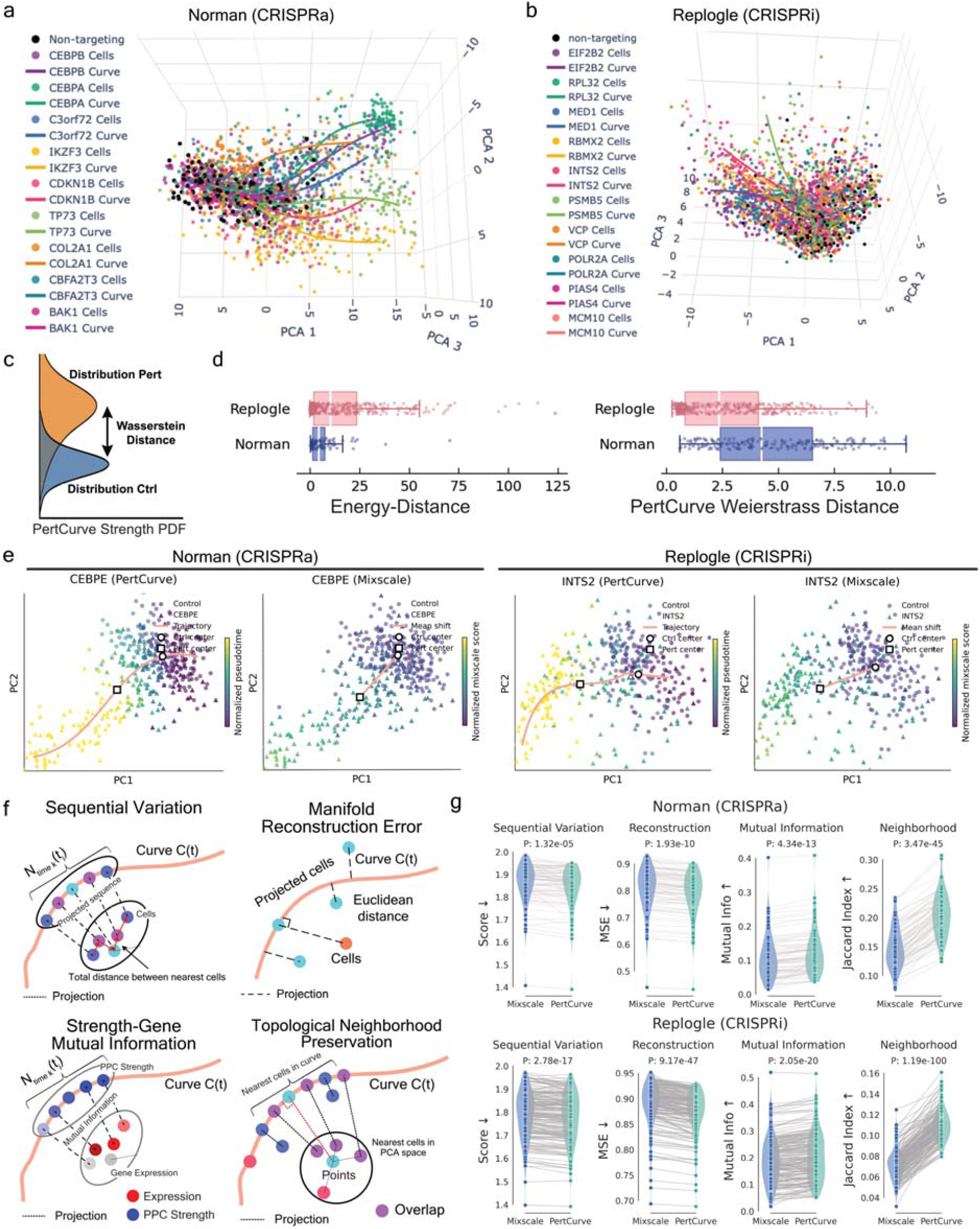
PertCurve improves upon prior methods by refining population- and single-cell level measurements. (a, b) Three-dimensional manifold illustrating PertCurve fits across diverse perturbations in the Norman (CRISPRa) and Replogle (CRISPRi) dataset. Ten highlighted genes denote principal elements identified via hierarchical clustering of the gene distance matrix. (c) Schematic of population-level metrics calculated by PertCurve. (d) Boxplots displaying population-level perturbation strengths calculated by scPerturb (*left*) and PertCurve (*right*) for both Norman (CRISPRa) and Replogle (CRISPRi) datasets. (e) Representative manifold scoring for *CEBPE* in the Norman (CRISPRa) dataset and *INTS2* in the Replogle (CRISPRi) dataset, comparing PertCurve and Mixscale. (f) Schematic illustrating the metrics for comparing PertCurve and Mixscale, including sequential variation, Manifold Reconstruction Error, Strength-Gene Mutual Information and Topological Neighborhood Preservation. (g) Benchmarking between PertCurve and Mixscale across the comparison metrics in both the Norman (CRISPRa) and Replogle (CRISPRi) datasets.

Next, we leveraged the PertCurve to quantify individual-level heterogeneity—variations in transcriptomic response trajectories among cells within the same perturbation condition. The position of each cell’s projection along the fitted curve serves as a proxy for its individual response strength. To standardize this measure, we defined the perturbation strength at the curve’s starting point as 0 (representing the unperturbed state) and at the endpoint as 1 (representing maximal perturbation), thereby scaling all cellular responses to the interval [0, 1]. To validate our approach, we compared the single-cell perturbation strength derived from PertCurve with that obtained from Mixscale^14^. The results demonstrate that our PertCurve approach more specifically conforms to the manifold structure generated by single gene CRISPR□based perturbations (**Fig. 2e**).

In addition, we compared PertCurve and Mixscale analyses using four classical trajectory-quality metrics: sequential variation, manifold reconstruction error, strength–gene mutual information, and topological neighborhood preservation, and evaluated their performance in producing informative, locally structure-preserving orderings. These metrics consistently show superior performance for the PertCurve, highlighting the advantage of our method (**Fig. 2f and 2g**).

An additional advantage of simultaneously capturing population-level and single-cell level response strength is that it enables the derivation of normalized population-level perturbation response strength. Conventional metrics for population-level perturbation strength fail to account for the actual expression changes of perturbed genes. In our framework, by ordering cells based on inferred perturbation strength, we reconstructed expression of target genes, the majority of which are biologically consistent (downregulated in CRISPRi or upregulated in CRISPRa) and are amenable to subsequent downstream experimental analysis (**Supplementary Fig. 2**). The expression change derived from reconstructed expression is then used to normalize measured population-level perturbation strength, producing a metric of perturbation effect that is comparable across all perturbed genes. (**Fig. 3a and 3b)**.

**Figure 3.**
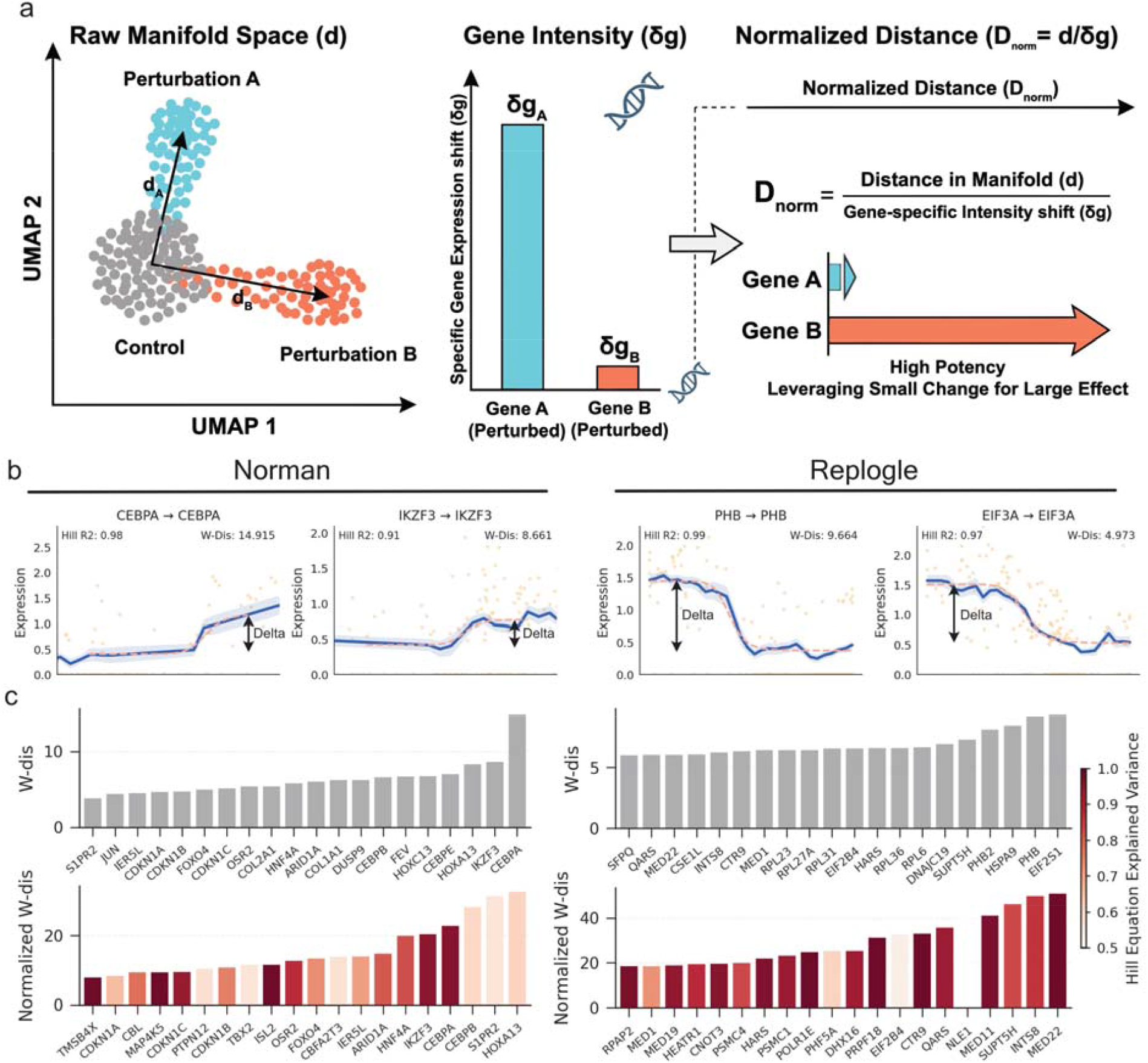
Normalized perturbation strengths prioritize genes with transcriptional regulatory functions. (a) Schematic comparison of absolute and normalized perturbation strengths. Conventional analyses emphasize global transcriptomic shifts but often overlook the intrinsic potency of a perturbation on its direct target. (b) PertCurve fitting of the expression trajectory of the perturbed target gene along the perturbation strength gradient, increasing with strength for CRISPRa (Norman) and decreasing for CRISPRi (Replogle). (c) Comparative ranking of genes by absolute and normalized perturbation effects. The top row displays genes with maximal absolute effects, predominantly functional genes, whereas the bottom row is sorted by normalized strength, prioritizing transcriptional regulatory genes. Colors by the Hill-equation R^2 as a measure of confidence.

PertCurve’s perturbation strength normalization reveals consistent patterns in both CRISPRa and CRISPRi experiments. In the Norman (CRISPRa) dataset, after normalization, the functional landscape centers on transcriptional regulation and epigenetic control (**Fig. 3c**). Key regulators involved in cell identity determination, chromatin remodeling, and cell-cycle progression—such as *HNF4A, ARID1A, FOXO4*, and *CDKN1C* and *CDKN1B*—rose to higher priority. *HNF4A* is a master transcription factor governing the identity of hepatocytes and pancreatic β-cells, regulating genes involved in glucose and lipid metabolism as well as hepatic detoxification^18^. *ARID1A*, a core subunit of the SWI/SNF chromatin remodeling complex, modulates chromatin accessibility to influence transcription initiation and DNA repair processes^19^. *FOXO4* mediates cellular stress responses by regulating cell cycle progression and apoptosis pathways^20^. *CDKN1C* and *CDKN1B* encode the cyclin-dependent kinase inhibitors p57 and p27, respectively, which function as critical negative regulators of cell cycle progression^21^. In contrast, before normalization, while some transcription factors were retained, their ranks were lower and overall importance diminished. Instead, the functional emphasis shifted toward tissue construction and terminal differentiation, such as genes driving extracellular matrix formation (*COL1A1, COL2A1*)^22^.

In the Replogle (CRISPRi) dataset, normalization similarly prioritized genes with core roles in transcriptional regulation and RNA metabolism (**Fig. 3c**). These include genes involved in the mediator complex for transcription initiation (*MED22, MED11*)^23^, key components that maintain transcriptional elongation (*SUPT5H, CTR9*)^24,25^, and subunit of Integrator complex and spliceosome (*INTS8, PRPF18*)^26^. Components of the Integrator complex and spliceosome ensure proper RNA maturation, highlighting the functional importance of coupled transcription-processing mechanisms. Conversely, the pre-normalization ranking was more biased toward ribosomal proteins (*RPL6, RPL36, RPL23*)^27^ and mitochondrial homeostasis factors (*PHB, PHB2, HSPA9*)^28^, which prioritizes protein production execution and cellular energy supply (**Fig. 3c**). Analyses in both datasets highlight the elevated priority of genes involved in transcriptional regulation after normalization, consistent with previous studies showing that knockdown of core genes involved in transcription initiation, mRNA synthesis, and chromatin remodeling significantly affects global RNA synthesis rates^29^, underscoring the biological relevance of normalized perturbation strength.

### PertCurve reveals response trajectory patterns and the underlying mechanism

For perturbation responses, conventional differential expression (DE) analysis only identifies genes with significant differences in average expression between perturbed and control conditions. Owing to cellular-level heterogeneity, DE analysis is prone to missing perturbation-responsive genes and cannot resolve dynamic response patterns of responsive genes across gradual perturbation strength, which are critical to understanding complex diseases and traits^15–17^. Previous dose–response studies have demonstrated that downstream target genes exhibit non-linear responses to perturbation or overexpression of transcription factors^15–17^, rendering linear models that do not account for cellular heterogeneity inadequate. The non-linear PertCurve method enables a more comprehensive and nuanced exploration of downstream transcriptional responses, and can resolve how downstream genes respond dynamically across the full continuum of perturbation strengths. Across both the Replogle (CRISPRi) and Norman (CRISPRa) datasets, we identified two overarching properties of downstream responses, modularity and asynchrony (**Fig. 4a and 4b**). Modularity refers to groups of genes exhibiting similar response patterns (co-regulated modules), while asynchrony indicates that different modules activate at distinct perturbation thresholds or follow divergent kinetics. From heatmaps of downstream gene expression ordered along perturbation strength and corresponding UMAP embeddings, it is clear that genes sharing similar expression trajectories cluster together to form several distinct groups in both CRISPRi and CRISPRa datasets, and response dynamics are markedly heterogeneous across these clusters (**Fig. 4a and 4b**; **Supplementary Fig 3a and 3b**).

**Figure 4.**
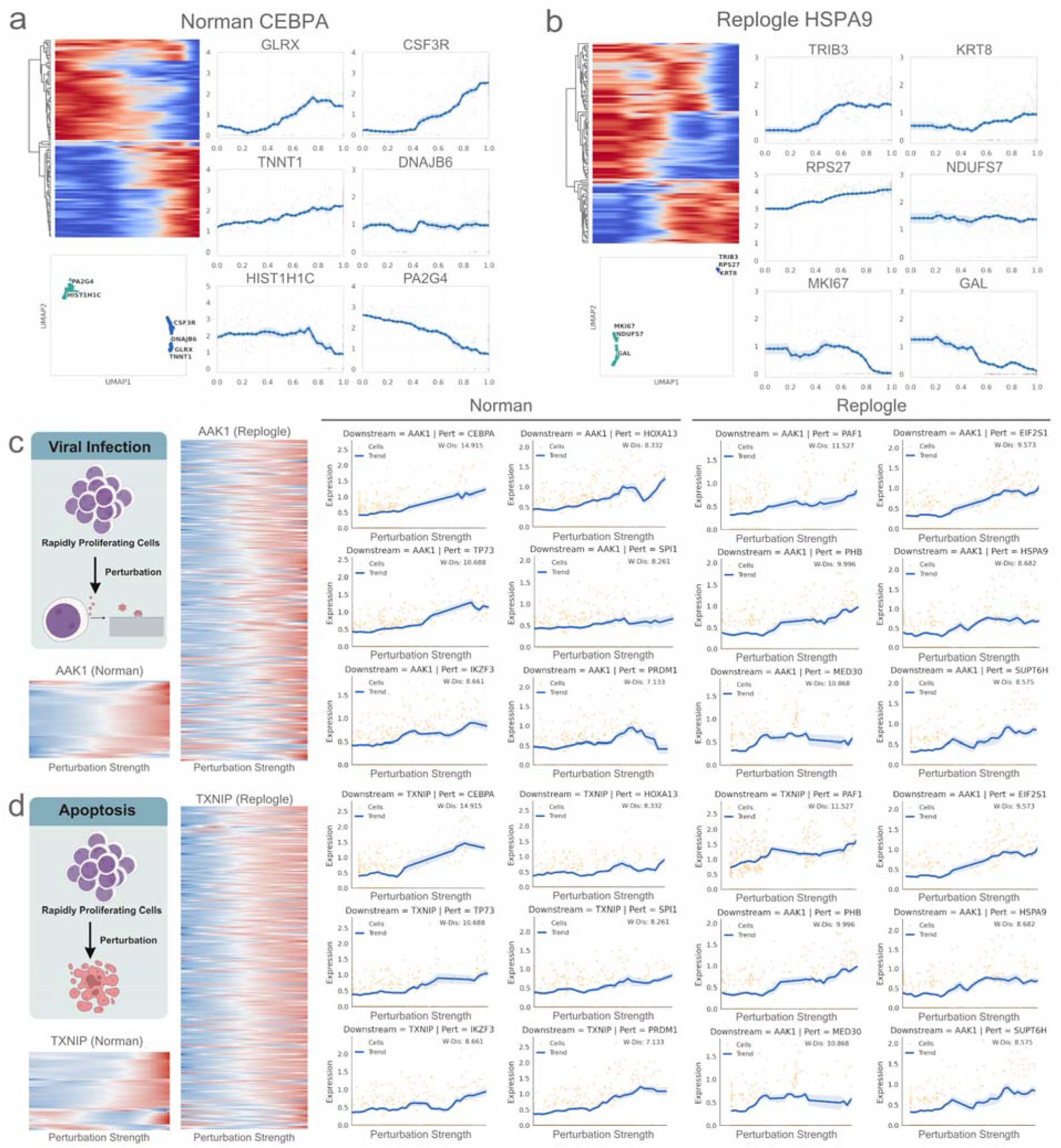
Downstream transcriptomic response trajectories reveal distinct response patterns, uncovering universal response patterns across CRISPRa and CRISPRi genetic modulations. (a, b) *Top left*, Normalized expression heatmaps of the top 100 differentially expressed downstream genes across a gradient of CRISPRa perturbation strengths targeting *CEBPA* in the Norman dataset (a) and CRISPRi perturbation strengths targeting *HSPA9* in the Replogle dataset (b). *Bottom left*, UMAP embeddings of downstream gene expression profiles under perturbations. Upregulated and downregulated genes segregate into two distinct clusters. *Right*, Example genes illustrating distinct downstream response dynamics for the perturbations. (c) Proportional-type up-regulation of viral-response and infection-mechanism genes: Perturbation triggers increased expression of genes associated with viral-entry pathways and innate immune signaling. The representative gene *AAK1* shows a pronounced increase, highlighting the cell’s systematic response to CRISPR-machinery delivery and presence. (d) Proportional-type up-regulation of apoptosis genes: Expression levels of genes involved in apoptosis increase post-perturbation. This is represented by *TXNIP*, which commonly increases upon CRISPR perturbations.

Furthermore, based on the previous descriptions of dynamic response patterns of responsive genes and our observations^13–17^, we have identified several recurrent response archetypes: proportional responses, in which expression scales linearly with perturbation strength; sensitive responses, where maximal changes occur at low perturbation levels; and threshold responses, in which expression changes only beyond a critical perturbation strength (**Fig. 4a and 4b**; **Supplementary Fig 3a and 3b**). These response archetypes recurred across the majority of perturbations in both the Replogle (CRISPRi) and Norman (CRISPRa) datasets. For examples, in the Norman (CRISPRa) dataset, under *CEBPA* perturbation, *TNNT1* exhibited a proportional increase, *PA2G4* a proportional decrease, *DNAJB6* remained largely unchanged, *HIST1H1C* displayed a threshold decrease, *CSF3R* showed a threshold increase, and *GLRX* demonstrated a sensitive increase (**Fig. 4a**). Similarly, under *HOXA13* perturbation (Norman CRISPRa), *GYPB* increased proportionally, *GAL* decreased proportionally, *NENF* remained stable, *CCNA2* exhibited a threshold decrease, *KRT8* a threshold increase, and *XIST* a sensitive increase (**Supplementary Fig 3a**). In the Replogle (CRISPRi) dataset, under *HSPA9* perturbation, *GAL* decreased proportionally, *RPS27* increased proportionally, *NDUFS7* was largely unaltered, *MKI67* exhibited a threshold decrease, *KRT8* a threshold increase, and *TRIB3* a sensitive increase (**Fig. 4b**). Correspondingly, under *EIF2S1* perturbation, *CCPG1* increased proportionally, *CKB* decreased proportionally, *KMT2A* was mostly unchanged, *CCNA2* showed a threshold decrease, *MALAT1* a threshold increase, and *TRIB3* a sensitive increase (**Supplementary Fig 3b**). These patterns were consistently reproduced across other individual perturbations.

More importantly, by integrating CRISPRi (Replogle) and CRISPRa (Norman) data from the same K562 cell line, which is a human chronic myeloid leukemia (CML) line carrying the BCR-ABL fusion oncogene due to the t(9;22) Philadelphia chromosome^30^, we uncovered universal response programs that are observed across most perturbations and are independent of the direction of perturbation (inhibition versus activation). For example, across most perturbations, genes associated with cell proliferation (*e*.*g*., *MKI67, CCNA2, CCNB1/2, CDK1, PLK1, CDC20, PTTG1, CDCA8, TOP2A, UB E2S, CDC45, RRM2*)^31^ consistently exhibited a threshold-type response: their expression remained stable at low perturbation strength but sharply declined to near zero once a critical threshold was surpassed, regardless of whether the upstream perturbation was inhibitory (CRISPRi) or activating (CRISPRa). This suggests a common “fail-safe” mechanism wherein sufficiently strong perturbations of any kind trigger cell cycle arrest in K562 (**Supplementary Fig. 4, Supplementary Fig. 5 and Supplementary Fig. 6**). Conversely, genes associated with apoptosis (*CASP8, DDIT3, TXNIP, TAX1BP1, BTG1/2, SOSTM1*)^32–35^ or viral infection response (*TRIM56, UBE2L6, MAP1LC3B, AAK1, SOCS1/2, PARP14*)^36–40^ displayed proportional-type upregulation: their expression increased steadily with perturbation strength across nearly all conditions. This implies that stronger perturbations, whether via repression or activation, will induce cumulative cellular stress, culminating in apoptotic priming and antiviral signaling (**Fig. 4c and 4d; Supplementary Fig. 5, Supplementary Fig. 7, and Supplementary Fig. 8**).

Beyond these universal response programs, downstream gene dynamics also revealed lineage-specific differentiation signatures. K562 cells possess multilineage potential and can be directed toward erythroid or myeloid fates (**Fig. 5a**)^4,5^. Our analysis demonstrates that the erythroid marker gene *HBA1* and the myeloid marker gene *AIF1* exhibited distinct expression trends under different perturbations. In the perturbations targeting *IKZF1, TP73*, and *PTPN1, HBA1* expression increased with perturbation strength, whereas *AIF1* expression remained unchanged or gradually decreased. Conversely, perturbations targeting *SPI1, CEBPA*, and *COL1A1* induced opposite patterns: *HBA1* expression remained stable or decreased, while *AIF1* expression increased (**Fig. 5b**). This reciprocal expression dynamic between *HBA1* and *AIF1* was further clarified in UMAP projections, where opposing trends distinctly separated different perturbations in both the Norman and Replogle datasets (**Fig. 5c**). Comparable lineage-biasing patterns were also observed for other erythroid marker genes, such as *HBZ* and *GYPA*, and myeloid marker genes, including *CTSC, IGFBP2, MS4A3* and *S100A11* (**Fig. 5d**). In contrast, the cell cycle-related genes, apoptosis-related genes and virus-related genes exhibit predominantly uniform directional changes, being consistently either upregulated or downregulated across perturbations, a pattern distinctly different from that observed for differentiation-associated genes (**Supplementary Fig. 5**).

**Figure 5.**
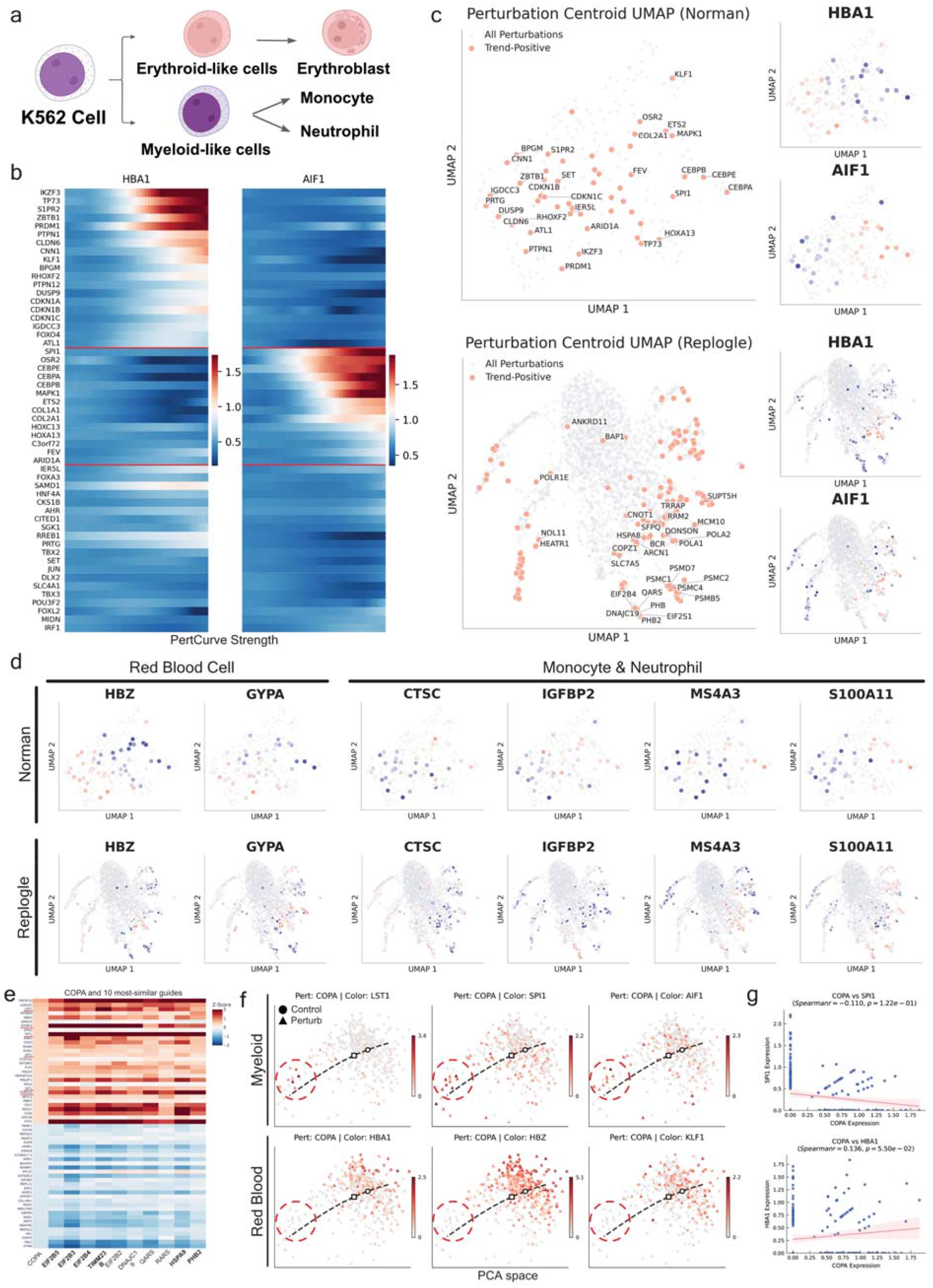
Downstream transcriptomic response trajectories delineate differentiation trends and identify masked context-specific genes. (a) Schematic of K562 multipotency and differentiation. (b) UMAP projection of perturbation-specific transcriptional centroids derived from the Norman and Replogle CRISPR-screening datasets. The embedding is annotated by target genes and colored by differential expression of lineage-specific markers, revealing how genetic interventions shift cellular states. (c) Divergent expression patterns of lineage markers under varying perturbation strengths. Expression trends of *HBA1* (erythroid marker) and *AIF1*(myeloid marker) are shown across a gradient of perturbation strength, indicating that perturbation strength differentially modulates lineage-specific transcriptional programs. (d) Functional landscape of perturbation-induced phenotypes. UMAP visualization of perturbation centroids colored by signature scores for erythroid and myeloid modules. Perturbations cluster according to their lineage-biasing effects, with a distinct subset of genes driving a transition toward systemic apoptotic signatures. (e) Representative *COPA* perturbation trajectory in the Replogle dataset. Cells are shown in PCA space and colored by expression of classical myeloid-associated genes (*LST1, SPI1* and *AIF1*) or erythroid-associated genes (*HBA1, HBG1* and *KLF1*). The fitted PertCurve trajectory orders cells from control-like states to high-strength perturbation states, revealing increased myeloid marker expression and reduced erythroid marker expression with increasing *COPA* perturbation strength. (f) Association between *COPA* perturbation strength and lineage-specific transcriptional programs. PertCurve-inferred perturbation strength is positively associated with the myeloid program and negatively associated with the erythroid program, indicating that high-strength *COPA* perturbation shifts K562 cells toward a myeloid-like state while suppressing erythroid differentiation. (g) Heatmap of top DEG and closest 10 perturbation for *COPA* perturbation from Genome-Wide Perturb-Seq website of Whitehead Institute. Y-axis label is manually labeled for format.

Furthermore, PertCurve can uncover differentiation signals masked by conventional methods. Conventional differential expression analysis is readily confounded by numerous low-perturbation-strength cells, obscuring true differentiation signals in high-perturbation-strength cells. Our PertCurve method stratifies cells by perturbation strength, clearly revealing the effects of high perturbation strength on cell differentiation. For instance, conventional differential expression-based methods only yield weak signals for *COPA* perturbation, as even the absolute z-scores of the top 30 upregulated and downregulated genes are all relatively low (**Fig. 5g**). By modeling cell-level heterogeneity through curve fitting, we recovered differentiation signals masked in conventional analyses. Myeloid markers *LST1, SPI1* and *AIF1* are upregulated in high-perturbation cells and downregulated in control and low-perturbation cells. Conversely, erythroid differentiation markers *HBA1, HBG1* and *KLF1* are highly expressed in control and low-perturbation cells, but lowly expressed in high *COPA* perturbation cells. Thus, high *COPA* perturbation promotes myeloid differentiation in K562 cells (**Fig. 5e and 5f**). Previous studies have established that pathogenic *COPA* mutations drive persistent retention of stimulator of interferon genes (STING) at the Golgi, resulting in constitutive STING activation^41^. Sustained STING signaling in turn rewires the lineage fate commitment of hematopoietic stem cells (HSCs), skewing their differentiation toward the myeloid lineage at the expense of B lymphopoiesis and erythropoiesis^42^. These findings are consistent with and support the conclusions of our study.

### PertCurve improves perturbation-response prediction and enables strength-aware evaluation

PertCurve-derived metrics can be further leveraged to refine perturbation predictive modeling and its evaluation. To demonstrate its utility, we integrated calculated cell-wise perturbation strength scores into GEARS, a state-of-the-art model for predicting transcriptional responses to genetic perturbations^43^.

In rigorous benchmarking experiments, the integration of PertCurve metrics consistently improved model performance. Evaluated across four distinct test splits—including unseen single-gene perturbations and novel double-gene combinations—PertCurve-enhanced GEARS model significantly outperformed the original version (**Fig. 6a**, *left*). This performance gain was quantitatively validated by improvements in both Mean Squared Error and Pearson correlation coefficients, assessed over the entire transcriptome as well as specifically over differentially expressed (DE) downstream gene subsets. Detailed case studies, such as perturbations targeting *TP73* and *PTPN1* within the test split, further illustrate these improvements (**Fig. 6a**, *right*).

**Figure 6.**
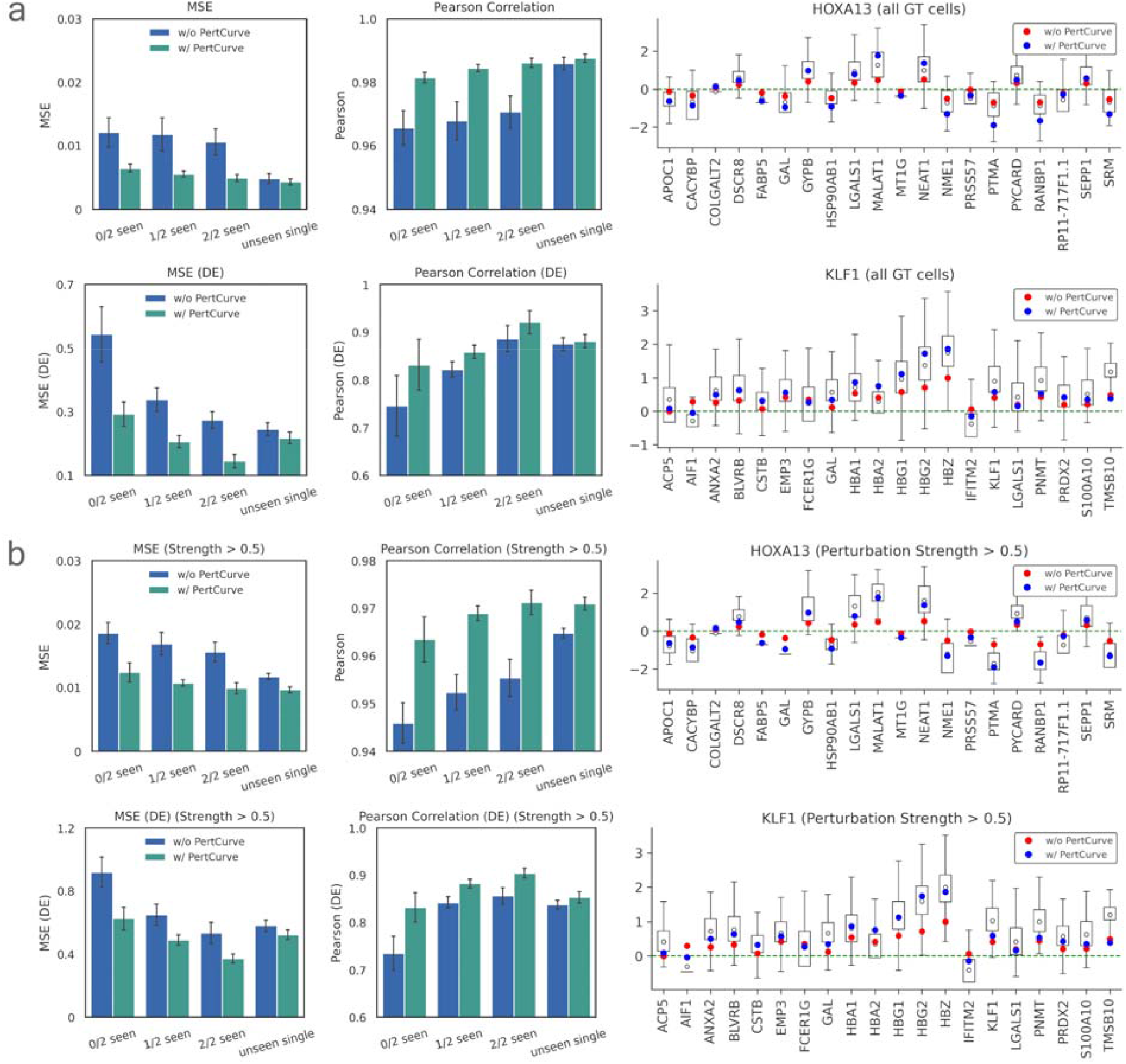
Integration of PertCurve metrics enhances predictive performance and biological fidelity in transcriptional response modeling. *(a) Left*: Global performance benchmarking of the original GEARS model and the PertCurve-enhanced GEARS model across four test types using all perturbed cells. Mean squared error (MSE) and Pearson correlation are evaluated over the full transcriptome and over differentially expressed (DE) downstream gene subsets. *Right*: Gene-level prediction fidelity for *TP73* and *PTPN1* perturbations. Boxplots show ground-truth single-cell expression distributions for top20 DE genes (alphabetically sorted), with deterministic model predictions overlaid for the original GEARS model (*red*) and the PertCurve-enhanced model (*blue*). (b) *Left*: Performance on high-strength perturbation cells, defined by perturbation score > 0.5. MSE and Pearson correlation are reported for both whole-transcriptome and differentially expressed (DE) gene subsets, highlighting model behavior on cells with stronger perturbation responses. *Right*: Gene-level prediction fidelity for *TP73* and *PTPN1* perturbations. Boxplots show ground-truth single-cell expression distributions for top20 DE genes (alphabetically sorted), with deterministic model predictions overlaid for the original GEARS model (*red*) and the PertCurve-enhanced model (*blue*).

Besides, current perturbation-prediction benchmarks typically define the ground-truth response by aggregating all cells assigned to a given perturbation. However, guide assignment does not necessarily imply effective perturbation, as labeled cell populations can contain weakly perturbed or escaper-like cells whose transcriptional states remain close to the unperturbed baseline.

We next asked whether PertCurve could provide a more biologically faithful evaluation target. To validate inference performance in detail, we compared the model’s point predictions against cells exhibiting high perturbation strength (**Fig. 6b, Supplementary Fig. 9**). Under both evaluation settings, the PertCurve-enhanced GEARS model consistently outperformed the original GEARS. This demonstrates that PertCurve-guided predictions more accurately capture the expression profiles of successfully perturbed cells, proving that incorporating continuous biological priors enhances predictive fidelity. Furthermore, a global evaluation across all perturbations shows that PertCurve consistently improves performance for low-sample perturbations situated at the tail of the distribution (**Supplementary Fig. 10**), establishing it as both an interpretable and robust framework for model enhancement. These results show PertCurve aids perturbation prediction by offering continuous biological priors for better model training and enabling strength-aware benchmarking that refines evaluation from a heterogeneous population to cells with clear perturbation responses.

## Discussion

Perturb-seq experiments enable large-scale profiling of single-cell responses to genetic interventions, providing a robust foundation for deciphering causal relationships within biological systems. However, the resulting data exhibit complex and multilayered heterogeneity that current analytical frameworks struggle to delineate and limiting the use of Perturb-seq to obtain biological insights. To address this, we introduce PertCurve, a method designed to simultaneously characterize perturbation-induced heterogeneity at both the population level and the single-cell level, and reveal biological insights that were previously unattainable.

Under various measurement metrics, PertCurve outperform the previously established scPerturb at the population level (across multiple cell groups) and surpass mixscale at the single-cell level. More importantly, by jointly analyzing heterogeneity at the population level and the single-cell level, our approach yields new insights into cellular perturbation responses. Notably, conventional tools often estimate perturbation strength, defined as the overall impact of modulating a target gene’s expression on the transcriptome, without explicitly accounting for the change in the target gene’s expression level. In contrast, our framework simultaneously quantifies perturbation strength and reconstructs the expression trajectory of the perturbed gene, thereby enabling derivation of a normalized perturbation strength. Normalization shifts the prioritization of perturbation effects from highly expressed structural genes toward core functional regulators governing cell identity and homeostasis, including transcription factors, chromatin remodelers, and RNA-processing components, offering a biologically grounded interpretation of perturbation outcomes. Studies on nascent transcriptome after perturbation show that knockdown of core genes involved in transcription initiation, mRNA synthesis and chromatin remodeling significantly affects global RNA synthesis rates, which supports the conclusions we presented above^29^.

Furthermore, PertCurve enables the discovery of previously overlooked patterns in downstream perturbation responses. Through cross-modal integration of CRISPRi and CRISPRa datasets, our method reveals the modular, asynchronous nature of transcriptional responses to single-gene perturbations. Extending prior experimental and computational findings^13–17^, it clusters these responses into recurrent archetypes. In K562 cells, we identify shared downstream response programs across diverse perturbations, irrespective of modality. For instance, genes associated with viral infection or apoptosis consistently exhibit a proportional increase in expression with increasing perturbation strength, whereas proliferation-related genes display threshold-dependent behavior, remaining stable until a critical threshold is reached, after which their expression declines sharply. The identification of consistent downstream response patterns across both CRISPRi and CRISPRa perturbations indicates that these changes are likely CRISPR-experiment-dependent, warranting consideration in future interpretation of Perturb-seq data.

Beyond these universal response programs, in context-specific responses, PertCurve enhances analytical robustness by filtering stochastic noise associated with off-target effects. In *COPA* perturbation experiments, failing to stratify cells by perturbation strength causes high-perturbation signals to be obscured by low-perturbation noise. Our method separates cells by perturbation strength, and clearly demonstrates that myeloid markers are upregulated while erythroid differentiation markers are downregulated in high *COPA* perturbation cells. This confirms that high-intensity *COPA* perturbation drives K562 cells toward myeloid differentiation, consistent with previous findings.

Current perturbation prediction models typically treat perturbations as binary states (perturbed vs. unperturbed) during model training. However, multiple studies have shown that even highly complex models often fail to outperform simple linear baselines, suggesting that increased model complexity does not necessarily translate into improved performance^44^. This limitation is potentially attributed to the oversimplified binary representation of perturbations, which fails to reflect biological reality. Several recent studies have proposed integrating normalized, dose-like perturbation strength as continuous inputs to improve model predictive power ^44^. In our experiment, integrating PertCurve-derived results, which capture heterogeneity in perturbation strength across cells under the same perturbation, into the state-of-the-art prediction model GEARs significantly improved its performance on prediction tasks, demonstrating the critical importance of accounting for cellular heterogeneity in predictive modeling. Meanwhile, heterogeneity in cellular transcriptional responses to identical genetic perturbations means that including low-perturbation-strength cells in the evaluation set compromises assessment performance. We therefore propose using only the successfully perturbed, *i*.*e*., top 50% of cells by perturbation strength, in the evaluation set as the metric for prediction model performance. Furthermore, our findings on the universal responses of certain downstream genes across diverse perturbations—even across opposing modalities such as repression (CRISPRi) and activation (CRISPRa)—indicate that the design of predictive model assessments may need to be revised. Specifically, such models could differentially handle genes with conserved response patterns versus those with perturbation-specific behaviors.

In summary, PertCurve is broadly applicable to the analysis of diverse perturbation assays, offers novel biological insights into previously overlooked perturbation responses, and provides actionable guidance for the development of the next generation of perturbation prediction models. Beyond CRISPR-based screens, it can also be extended to other contexts involving single-factor-induced global cellular changes, such as those induced by drug treatments.

## Methods

### Perturbation Curve Fitting

Let ***X*** ∈ *R*^*N*×*G*^ denote the single-cell gene expression matrix containing *N* cells and *G* genes. For a specific genetic perturbation target *p*, we define a perturbation-class indicator vector ***c*** ∈{0,1}^*N*^, where *c*_*i*_ = 0 denotes a cell in the wet-lab control state and *c*_*i*_ = 1 denotes a cell subjected to the wet-lab perturbation condition. Due to the inherent singlecell response observed in CRISPR interference or CRISPR activation screens, cells where *c*_*i*_ = 1 do not exhibit a uniform transcriptomic shift but rather a continuous gradient of perturbation efficacy. Our objective is to infer a continuous scalar variable *t*_*i*_ ∈ [0,1] for each cell to represent its precise perturbation strength, and to model the dynamic response of any downstream gene *g* of the perturbation *p* as a continuous function of *g*_*p*_ =*g*_*p*_ (*t*). To achieve our objective, we have carried out the following steps. First, single-cell CRISPR screening data often exhibits extreme class imbalance (*N*_*ctrl*_ » *N*_*pert*_) where the unperturbed control cells vastly outnumber the cells in any specific perturbation group. To mitigate this, we employed a subsampling strategy to equalize the density contribution of the control cells relative to the perturbed populations |*s*_*ctrl*_ |≈|s_*pert*_|. This setting follows the scperturb preprocessing procedure.

After preprocessing and reducing the dimensionality of the target and control cells *P* to via Principal Component Analysis (PCA) on their transcriptomic expression, we define the primary axis of variation as the vector connecting the control and perturbation centroids.

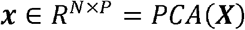

Then, we compute the geometric centroids of the control and perturbed populations in the principal component space.

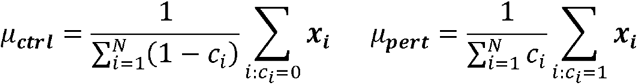

Initializing the fitting curve with a directional vector 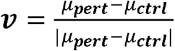 anchored to the control and perturbation centroids is critical because standard unsupervised manifold learning algorithms lack directionality. CRISPR screens inherently contain a large fraction of “escaper” cells that received a guide RNA but show no detectable phenotypic knockout, so the perturbed and control cell clouds often heavily overlap. Anchoring the vector mathematically cuts through this dense overlapping noise and it can successfully trace the true dose-response trajectory of the genuinely perturbed cells.

After this, our algorithm implements a “Binned Skeleton” strategy. We project all cells onto this one-dimensional axis to obtain a projection scalar *s*_*i*_ = (**x**_**i**_ − *μ*_**ctrl**_)^*T*^**v**. The projected space is uniformly partitioned into *K* discrete bins. Let ℬ_*h*_ denote the set of cells assigned to the k-th bin, we aggregate the imbalanced-density cells by calculating a set of *K* geometric centroid **a**_**k**_.

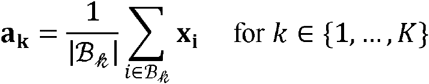

A smoothed curve, defined as a vector-valued function **f**(*s*) ∈ *R*^*G*^, is fitted through the ordered anchors {**a**_1_, …,a_**K**_} utilizing univariate splines. In practice, to ensure computational tractability and robustness, this continuous manifold is evaluated as a finely sampled sequence of *M* points ℱ = {**f**_1_, … **f**_M_}.

Finally, for each cell i, we determine its orthogonal projection onto the manifold by identifying the index *m*_*i*_ of the closest discrete curve point:

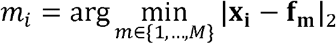

To accurately represent the cellular transition state while accounting for the varying geometry of the high-dimensional manifold, the raw score *τ*_*i*_ is computed as the cumulative geodesic arc length along the curve up to the projected index *m*_*i*_ where *τ*_*i*_ = 0 if *m*_*i*_ = 1.

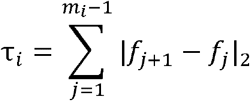

Then, the perturbation curve score *t*_*i*_ for each cell is obtained via min-max normalization of the cumulative arc lengths ensuring that *t*_i_ ∈ [0,1], where *t* = 0 strictly corresponds to the unperturbed baseline state and *t* = 1 represents the maximal perturbation efficacy observed in the dataset.

Curve fitting and cell projection are performed iteratively: after an initial trajectory is estimated, cells are re-binned according to their current projections, bin centroids are recomputed, and the spline curve is refitted until convergence or until a maximum number of iterations is reached. This joint refinement stabilizes the inferred manifold and yields the final continuous perturbation strength score for each cell.

### Quantitative Evaluation of Inferred Trajectories

To systematically benchmark the fidelity and robustness of the continuous perturbation manifolds inferred by PertCurve, we established a comprehensive set of quantitative evaluation metrics. These metrics assess the inferred strength *t* from multiple geometric and statistical perspectives, focusing specifically on highly informative differentially expressed genes to isolate the true perturbation signal from background biological noise.

#### Sequential Variation

Biological state transitions induced by perturbations are expected to manifest as continuous transcriptional gradients rather than erratic jumps. We evaluate the local variation of gene expression along the inferred strength trajectory. For cells ordered by their inferred score t, the variation is calculated as the ratio of the mean squared difference between adjacent cells to the total global variance:

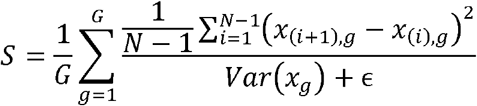

where *x*_(*i*),*g*_ represents the expression of gene g in the i-th cell along the sorted trajectory. A lower S score indicates a smoother, more biologically plausible transition.

#### Manifold Reconstruction Error

To quantify how well the 1D continuous variable explains the high-dimensional structural variance, we calculate the Mean Squared Error (MSE) between the standardized original expression matrix *x* and a reconstructed trajectory matrix 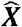 which is approximated via a local smoothing kernel (sliding window) along the ordered strength *t*. The reconstruction is approximated via a local convolution-based sliding window applied over the ordered cells.

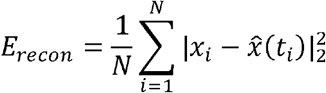

A lower reconstruction error implies that the strength axis successfully encapsulates the dominant biological variation induced by the targeted perturbation without severe information loss.

#### Pseudotime-Gene Mutual Information

Standard linear correlation coefficients often fail to capture the complex, non-linear dependencies inherent in downstream gene regulatory responses. To overcome this, we compute the average Mutual Information between the inferred strength (a kind of pseudo-time) *t* and the continuous expression profiles of top responsive genes.

The continuous MI is defined as:

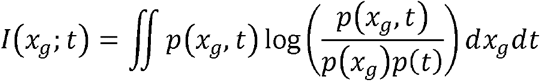

In practice, *I*(*x*_*g*_;*t*), is estimated utilizing k-nearest neighbors approach for continuous variables. To evaluate the trajectory’s global informational yield, we calculate the arithmetic mean of MI across a subset of top differentially expressed genes *g*_*Dℰ*_.

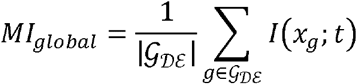

A higher global MI signifies that the inferred strength ordering possesses a higher informational density and accurately encodes the dynamic regulatory behavior of responsive targets.

#### Topological Neighborhood Preservation

A mathematically sound 1D reduction should preserve the intrinsic high-dimensional geometry of the cell cloud. For each cell, we identify its k-nearest neighbors in both the high-dimensional PCA space *N*_*HD*_ and the 1D strength space *N*_l*D*_. The topological fidelity is evaluated using the Jaccard index of these neighbor sets:

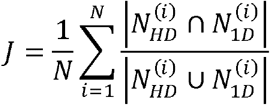

A higher Jaccard similarity indicates superior preservation of the genuine cellular micro-environments.

### Quantification of global perturbation magnitude

#### Projection-Distribution Metric: Wasserstein Distance by Optimal Transport

The Wasserstein distance (specifically the Wasserstein-1 or Earth Mover’s Distance) respects the underlying geometric topology of the transcriptomic manifold. It calculates the minimum global transport cost required to structurally transform the unperturbed baseline state into the perturbed state:

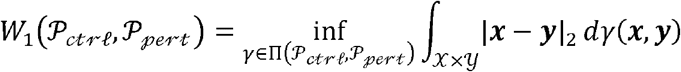

where Π(*P*_*ctrl*_,*P*_*pert*_) denotes the set of all joint probability measures γ(*x,y*)with marginals *P*_*ctrl*_ and *P*_*pert*:_.

#### Energy-distance baseline

As a baseline, we follow the established ***scperturb*** statistical framework designed for large-scale CRISPR screens and incorporate the Energy distance. The E-distance is a highly scalable, non-parametric metric that acts as a multivariate generalization of the squared difference of means, inherently adjusting for intra-population biological noise. It is defined only by PCA processed sample transcriptomics embeddings.

Let *x*_l_, …, *x*_*N*_ ∈ ℝ^*d*^ and *y*_l_, …, *y*_*N*_ ∈ ℝ^*d*^ be N and M cell samples from 2 distributions X and Y. E-distance metric is defined as below.

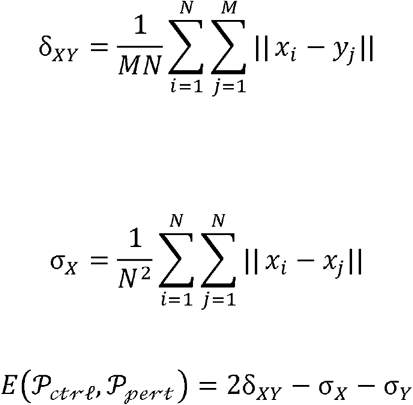

### Dynamic gene modeling via a Hurdle Mixture framework

While recent computational frameworks, such as Mixscale, have advanced the estimation of continuous perturbation efficacies at the single-cell level, their downstream evaluation relies predominantly on unimodal count-based generalized linear models, such as linear regression or Negative Binomial regression. Consequently, prior analyses have largely failed to accurately characterize the complex non-linear dynamic trends of downstream responsive genes along the graded strength of a given perturbation.

Unimodal regression approaches assume a single underlying distribution governed by a linear coefficient, inherently conflating the probability of a gene being actively transcribed with its absolute transcript abundance. Because single-cell RNA-seq data is fundamentally bimodal characterized by extreme sparsity and zero-inflation driven by both technical dropouts and genuine biological silencing (e.g., CRISPRi repression) can independently result in zero observed counts, it requires a specialized statistical treatment. To address this methodological gap and explicitly model the dynamic, non-linear responses of any downstream gene *g* along our inferred strength manifold *t*, we eschew standard unimodal regression in favor of a Hurdle-inspired mixture model.

This formulation implicitly decouples the binary transcriptional activation probability from the continuous expression magnitude. For a given local strength bin *ℬ*_*h*_, the expected expression of gene *g* (denoted as *E*[*y*_*ig*_]) is modeled as the product of the non-zero expectation and the probability of detection, where *α* is the sparsity hyperparameter set by properties of perturbation method.

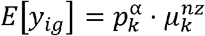

and the possibility and expectation is:

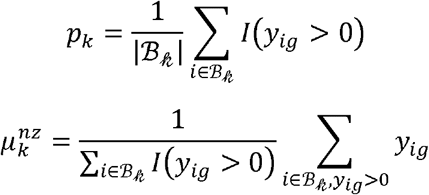

By decomposing the expectation, the model handles dropout artifacts and captures the true probability density transition of the cell state.

By decomposing the expectation, the model effectively accounts for dropout artifacts while capturing the genuine probability density transition of the cell state. This Hurdle-based framework thus enables robust characterization of downstream gene expression trends across varying perturbation strengths, regardless of whether the downstream gene is identical to or distinct from the targeted perturbation gene.

### Quantitative dose-response parameterization

To systematically characterize the topology of downstream gene responses relative to the perturbation gradient, we model the smoothed expression trend 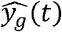 using a 4-parameter log-logistic (Hill) equation. This allows us to distill complex transcriptional shifts into interpretable dynamic parameters Θ_*g*_ = {*b*_*g*_,*c*_*g*_,*d*_*g*_,*e*_*g*_} learned for each responsive gene by minimizing the residual sum of squares.

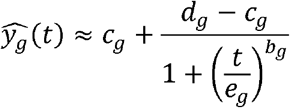

The loss function is defined as ℒ:

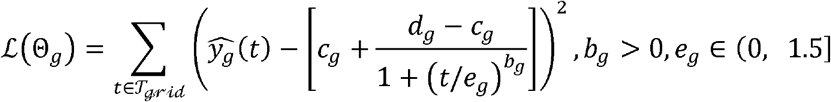

Here,*dg* captures the baseline expression at *t* = 0,*c*_*g*_ represents the asymptotic response limit as *t* → +∞,*e*_*g*_ designates the half-maximal effective perturbation strength (equivalent to the EC50) and *b*_*g*_ denotes the Hill coefficient. The *b*_*g*_ quantifies the non-linearity of the response, distinguishing between continuous, gradual shifts and abrupt, switch-like genetic responses to the upstream perturbation.

In addition, we can use the Hill-R^2^, defined as the variance explained by the fitted Hill equation on the downstream gene response curve, as a confidence metric to filter for top-responsive genes.

### Ranking Perturbation Sensitivity via Perturbation-level strength and individual-fitting Hill Dynamics

To systematically prioritize perturbations that exhibit significant distributional shifts relative to their modeled dynamic, we developed a composite ranking metric integrating perturbation divergence with non-linear curve fitting parameters. While Hill curve modeling characterizes the asymptotic dynamic range of a perturbation’s effect—defined by the differential between the upper and lower asymptotes (*Hill*_*c*_ and *Hill*_*d*_)—measurements of distributional shift such as wasserstein capture the total probabilistic deviation between control and perturbed states.

Standard differential expression approaches, which rely primarily on metrics such as log_2_ or absolute fold-change, are inherently limited to detecting shift significance in the arithmetic mean and are incomparable between different perturbations. Consequently, these methods are disproportionately biased toward highly-expressed genes and often may hindered from the labeled-but-untargeted CRISPR cells and technique dropout whose label are intrinsically unconfident.

In cell pathway multiplexed settings, true regulatory drivers often exhibit strict homeostatic buffering, meaning their absolute dynamic range may remain constrained but respond to perturbation by inducing complex shape-shifts in the population distribution, such as transitioning to bimodal states or amplifying transcriptional noise.

To isolate these non-linear drivers of cell fate, we sought to compute the ratio of the geometric state divergence to the gene’s asymptotic dynamic range, effectively measuring the topological sensitivity of the gene.

Conversely, the modeled dynamic range |*Hill*_*c*_ − *Hill*_*c*_| scales linearly with respect to transcriptomic amplitude. Dividing a quadratically growing numerator by a linearly growing denominator induces severe scale dependence, causing the metric to artificially inflate the importance of genes with extreme baseline variances.

We define the Normalized Perturbation Effect as:

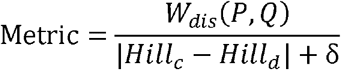

where *P* and *Q* represent the discretized probability distributions of the control and perturbed states along the normalized strength axis, and *δ* serves as a basal variance regularization term to prevent singularity artifacts (division by zero) in cases where Hill equation is near a line, or we direct filter them.

This normalization is mathematically justified because the Wasserstein distance and the Hill dynamic range operate on commensurate scales. Both metrics fundamentally reflect the magnitude of transcriptomic displacement—one capturing the global geometric divergence in the reduced manifold, the other capturing the gene-specific asymptotic expression amplitude along the inferred trajectory. Since the PCA embedding and the gene-wise smoothing operate on the same underlying expression matrix after identical preprocessing, the numerical outputs of two metrics are expressed in comparable dimensionless units of transcriptional variation.

By scaling the Wasserstein distance by the specific kinetic capacity of the gene, this formulation precisely quantifies the efficiency of the transcriptomic response. It actively prioritizes master regulators that execute profound, system-wide state reconfigurations with minimal metabolic expenditure.

### Technique details of incorporating PertCurve into GEARS

To inject trajectory information into GEARS, we first assigned each cell a perturbation score derived from PertCurve. For each perturbation condition, we still perform balanced downsampling to mitigate abundance biases. After fitting the curve, we re-projected all cells in the analysis set onto the fitted curve, computed their arc-length positions, and normalized these values into PertCurve score, with control cells anchored at zero. Since the classic GEARS model is trained on the whole processed Norman dataset including both single- and double-gene perturbations, we extended this PertCurve scoring workflow to two-gene combinatorial conditions to ensure a fair benchmark. While modeling dual-gene interactions via a single trajectory represents a simplified approximation of joint perturbation dynamics, it maintains structural uniformity across all benchmarking comparisons.

Formally, for each cell i, let *PC*_*i*_ denote its score calculated by PertCurve. This modulates the perturbation pathway during the forward pass of GEARS. Given the score, a learnable linear map produces a hidden-dimensional score vector, which is then converted into a residual gate:

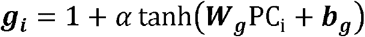

where *α*= 0.5 is a fixed gating amplitude.

Let *e*_*pert*_ denote the perturbation embedding after propagation on the perturbation graph, this modulates the global perturbation embedding and cells with higher PertCurve scores receive stronger perturbation signals while weakly perturbed or escaper cells receive attenuated signals:

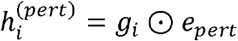

Thus, cells with higher PertCurve scores receive a stronger perturbation signal, whereas weakly perturbed or escaper-like cells receive an attenuated perturbation embedding. The gated perturbation representation is then fused with the base gene representation and decoded by the standard GEARS architecture.

### Evaluation protocol for prediction

We compared the original GEARS and a leakage-free PertCurve setting used for the main benchmark. During training of the score-aware model, PertCurve scores were attached to each training graph as cell-level inputs. During leakage-free evaluation, we assigned the same constant score (0.5) to every test graph, treating this scalar as a fixed model-design prior rather than test-time side information.

Therefore, the main comparison asks whether a score-aware GEARS architecture trained with PertCurve supervision remains beneficial with no strength leakage. Model performance was measured on the standard GEARS test split using mean squared error (MSE) and Pearson correlation, both across all genes and across the top differentially expressed downstream genes.

To ensure robustness against split-specific variance, all GEARS evaluations were repeated across multiple random seeds (1 to 10 by default), and the reported results were summarized across these runs.

## Statistics and reproducibility

Statistical analysis and data illustration was performed in python using the scanpy, numpy, torch, matplotlib.

The customized codes to perform the analyses described in this study are provided below (see ‘Code availability statement’). Same preprocessed method in scPerturb was used to determine sample size before algorithm. For random seeds, we apply split 1 to 10 in GEARS evaluation.

For the mixscale protocols, the official package is https://github.com/satijalab/Mixscale/. R package SeuratDisk was used to transform h5ad data to Seurat to be tested by mixscale. For scPerturb methods, we use both its official preprocess and energy-distance calculation script in https://github.com/sanderlab/scPerturb/. For the GEARS predictive method, the official package: https://github.com/snap-stanford/GEARS/.

## Author contributions

H.J. and X.Q. supervised the project. L.W., Y.Z., H.J. and X.Q. conceptualized the study and designed the experiments. Y.Z. implemented the code. L.W., Y.Z., H.J. and X.Q. performed the analysis and wrote the paper. G.Y., Y.L. provided assistance in writing and analysis.

## Acknowledgements

Acknowledgement This work was supported by the Hong Kong Research Grants Council General Research Fund (17102722, 17300523, 17302324) and the National Natural Science Foundation of China (32271445) to H.J. The work was conducted at the JC STEM Lab of Molecular Imaging, funded by The Hong Kong Jockey Club Charities Trust. For the purpose of Open Access, the author has applied a CC BY public copyright licence to any Author Accepted Manuscript version arising from this submission.

## Availability of data and materials

Norman dataset is available at https://www.ncbi.nlm.nih.gov/geo/query/acc.cgi?acc=GSE133344, and Replogle dataset can be downloaded at https://doi.org/10.25452/figshare.plus.20029387. Process script and processed data used for reproducibility are provided on Zenodo (DOI: 10.5281/zenodo.20643438).

The source codes of PertCurve as well as examples for testing are publicly accessible via https://github.com/yunhuazhong/PertCurve.

## Ethics declarations

The authors declare that they have no competing interests.

## Supplementary Figures

**Supplementary Figure 1.**
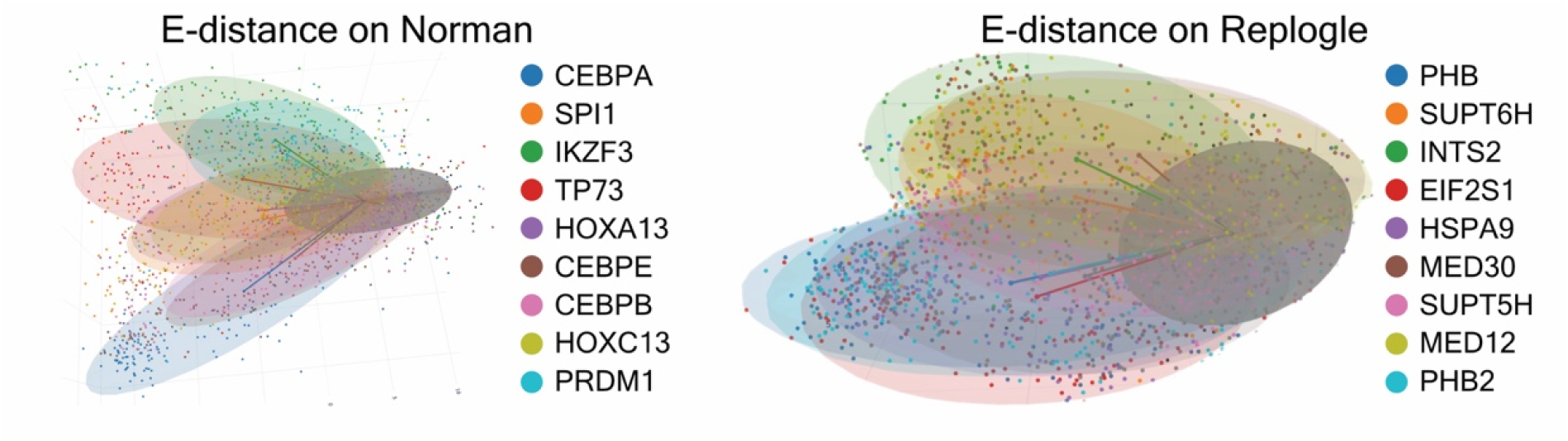
Three-dimensional PCA projection of the heterogeneity of cell states. Visualization of the top 10 most heterogeneous genetic perturbations and their E-distance visual illustrations from the Norman and Replogle datasets. Individual cells are embedded in a 3D PCA space and color-coded by their respective experimental perturbation conditions. Notably, E-distance is more sensitive to outlier cells.

**Supplementary Figure 2.**
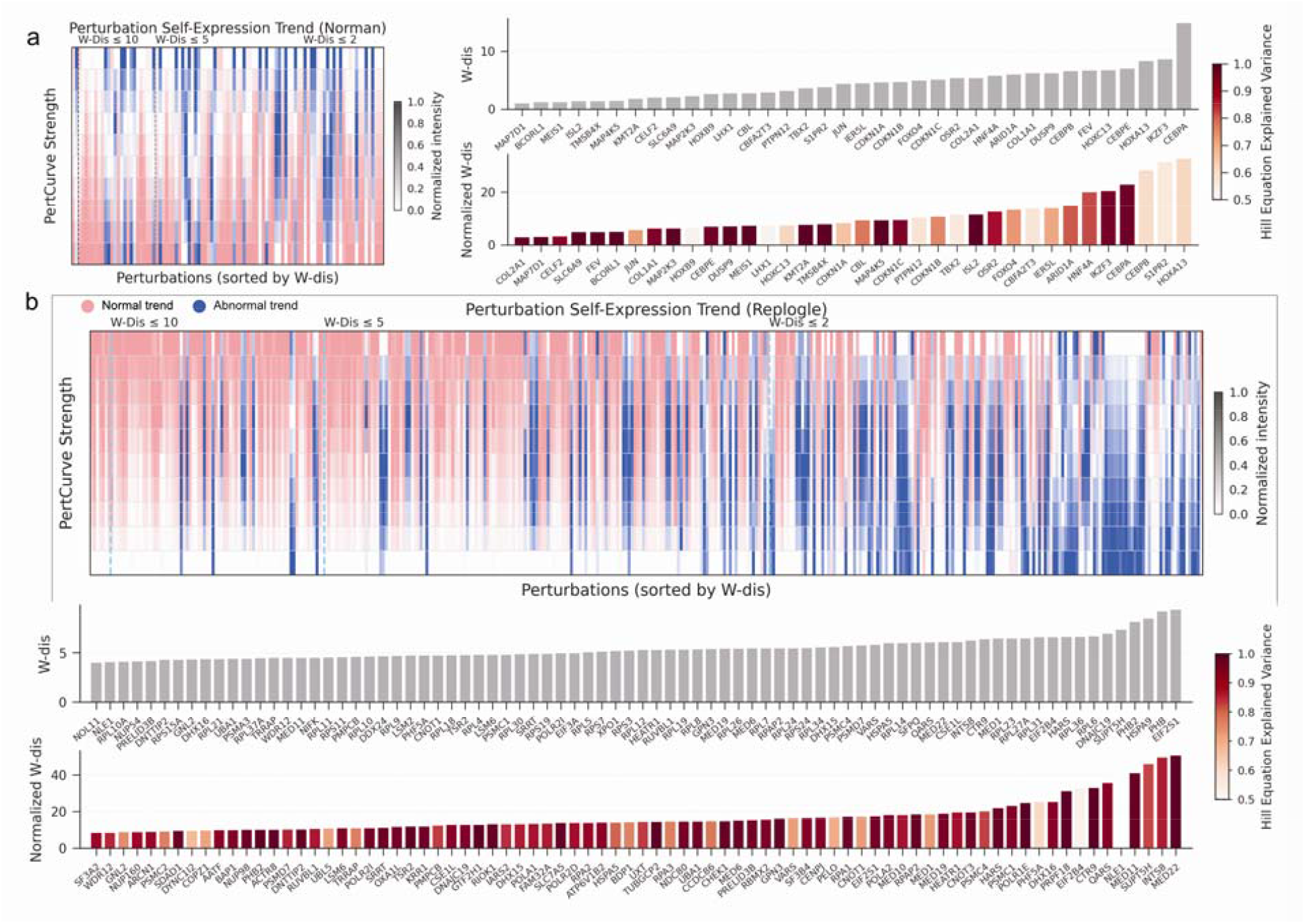
Reconstructed Target Gene Expression Trajectories & Extended Gene List of Normalized Perturbation Strengths. (a) Left: Recovery of perturbation□induced expression trends for target genes in the Norman. Most perturbations (pink) align with expected CRISPR dynamics (upregulation via CRISPRa) with higher□metric perturbations showing stronger adherence to these trends. Outliers (blue) may reflect technical constraints or novel biological properties. Right: Comparative ranking of genes (extended) by absolute and normalized perturbation effects in Norman dataset. The top row displays genes with maximal absolute effects, while the bottom row is sorted by normalized strength and colored by the Hill equation R^2^ as a measure of confidence. (b) Top: Recovery of perturbation□induced expression trends for target genes in the Replogle. Most perturbations (pink) align with expected CRISPR dynamics (downregulation via CRISPRi). Bottom: Comparative ranking of genes (extended) by absolute and normalized perturbation effects in Replogle dataset. The top row displays genes with maximal absolute effects, while the bottom row is sorted by normalized strength and colored by the Hill equation R^2^ as a measure of confidence

**Supplementary Figure 3.**
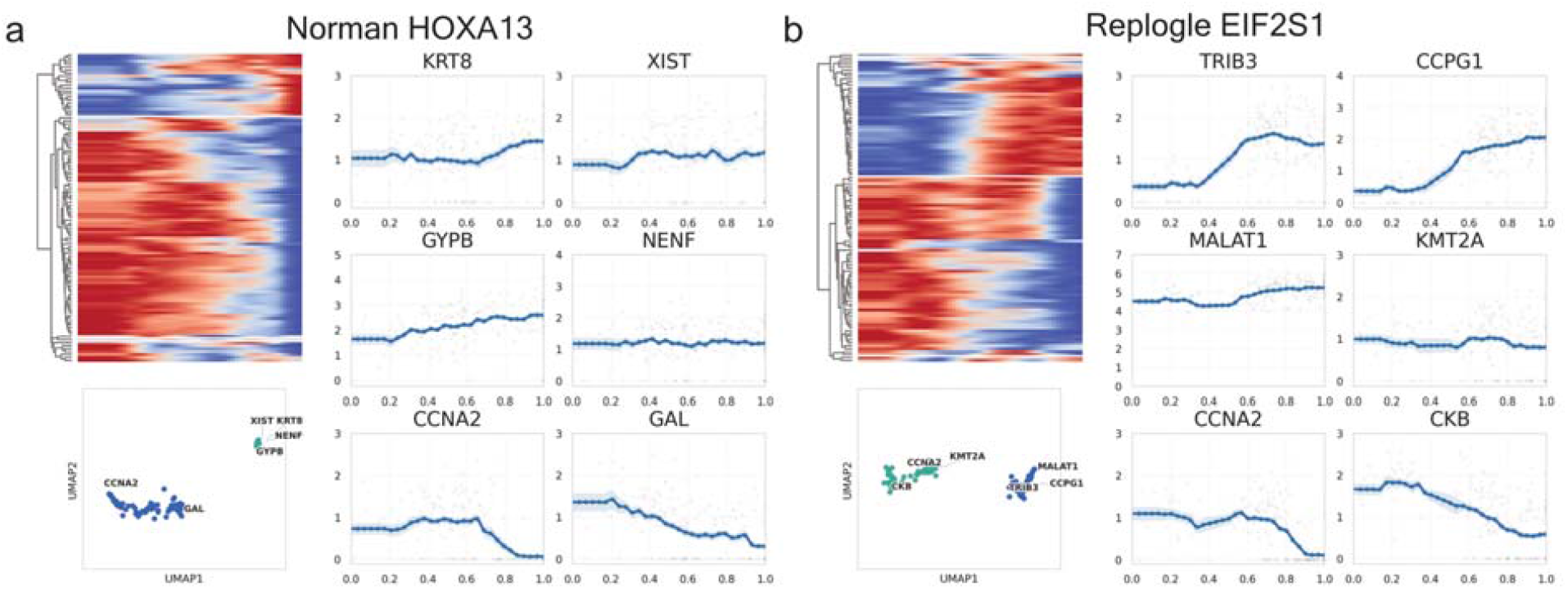
Downstream transcriptomic response trajectories exhibit multiple heterogeneous dynamic patterns that recur across distinct CRISPR perturbation modalities. (a, b). Top left, normalized expression heatmaps of the top 100 differentially expressed downstream genes across a gradient of CRISPRa perturbation strengths targeting *HOXA13* in the Norman dataset (a) and CRISPRi perturbation strengths targeting *EIF2S1* in the Replogle dataset (b). Bottom left, UMAP embeddings of downstream gene expression profiles under *HOXA13* perturbations (a) and *EIF2S1* perturbations (b), where upregulated and downregulated genes separate into two distinct clusters. Right: Example genes illustrating distinct downstream response dynamics for *HOXA13* (a) and *EIF2S1* (b) perturbations.

**Supplementary Figure 4.**
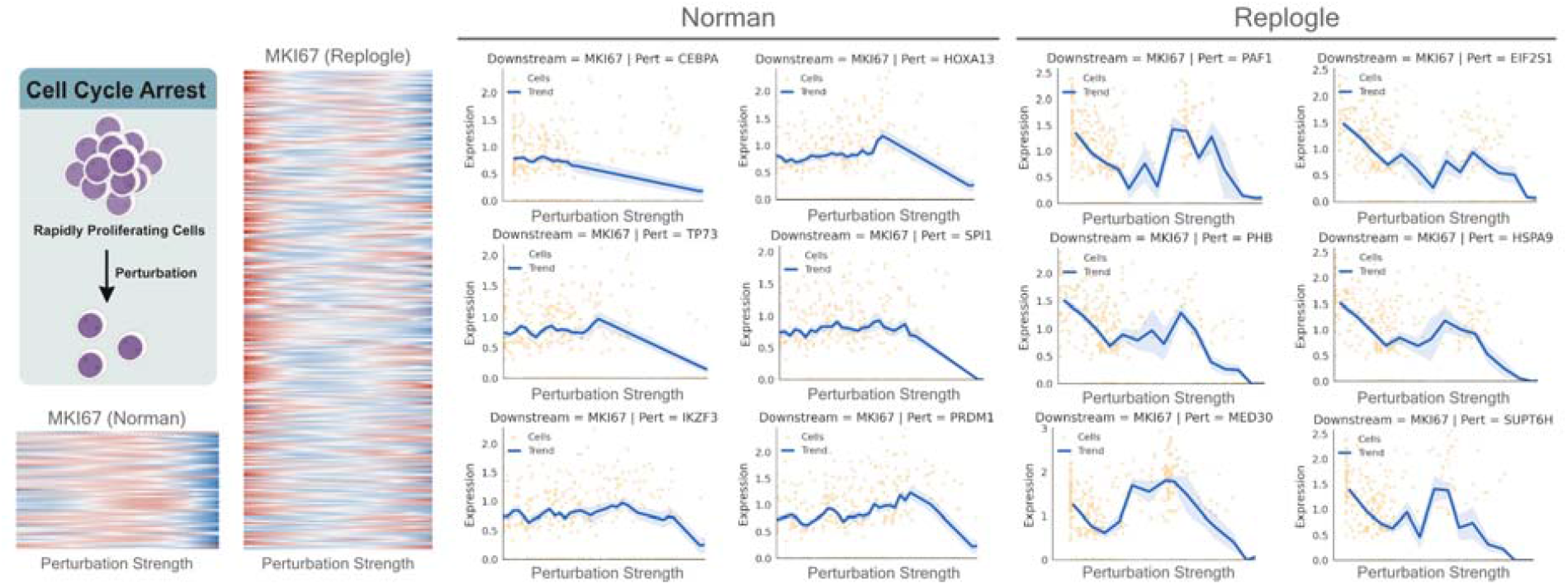
Universal response patterns on cell-proliferation genes across CRISPRa and CRISPRi genetic modulations. Threshold-type response of cell-proliferation genes: their expression remained stable at low perturbation strength but sharply declined to near zero once a critical threshold was surpassed. The representative gene *MKI67* exemplifies this trend, indicating a generalized suppression of the mitotic cycle.

**Supplementary Figure 5.**
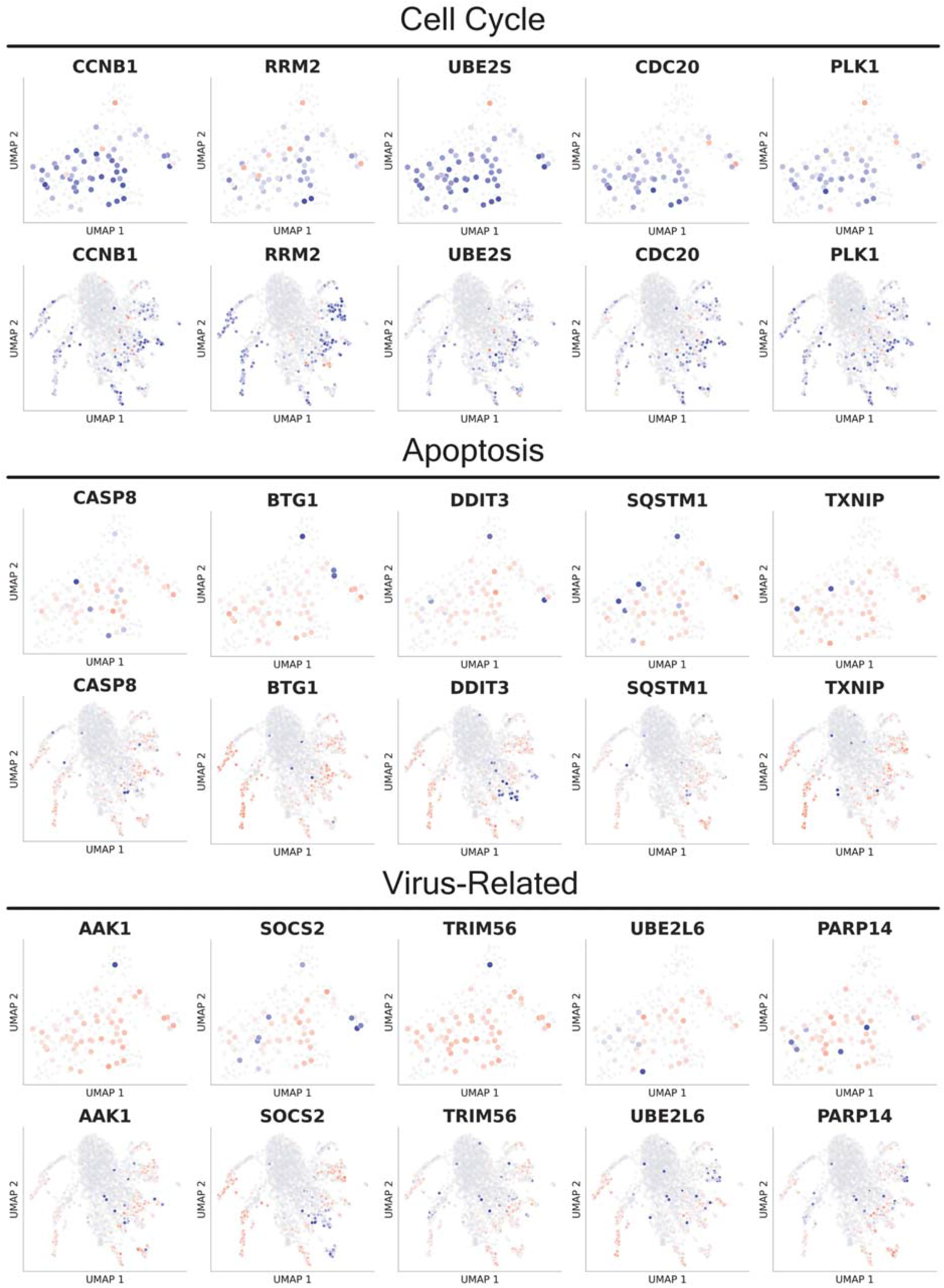
Functional landscape of perturbation-induced phenotypes. UMAP visualization of perturbation centroids colored by signature scores for apoptosis, viral infection and cell cycle.

**Supplementary Figure 6.**
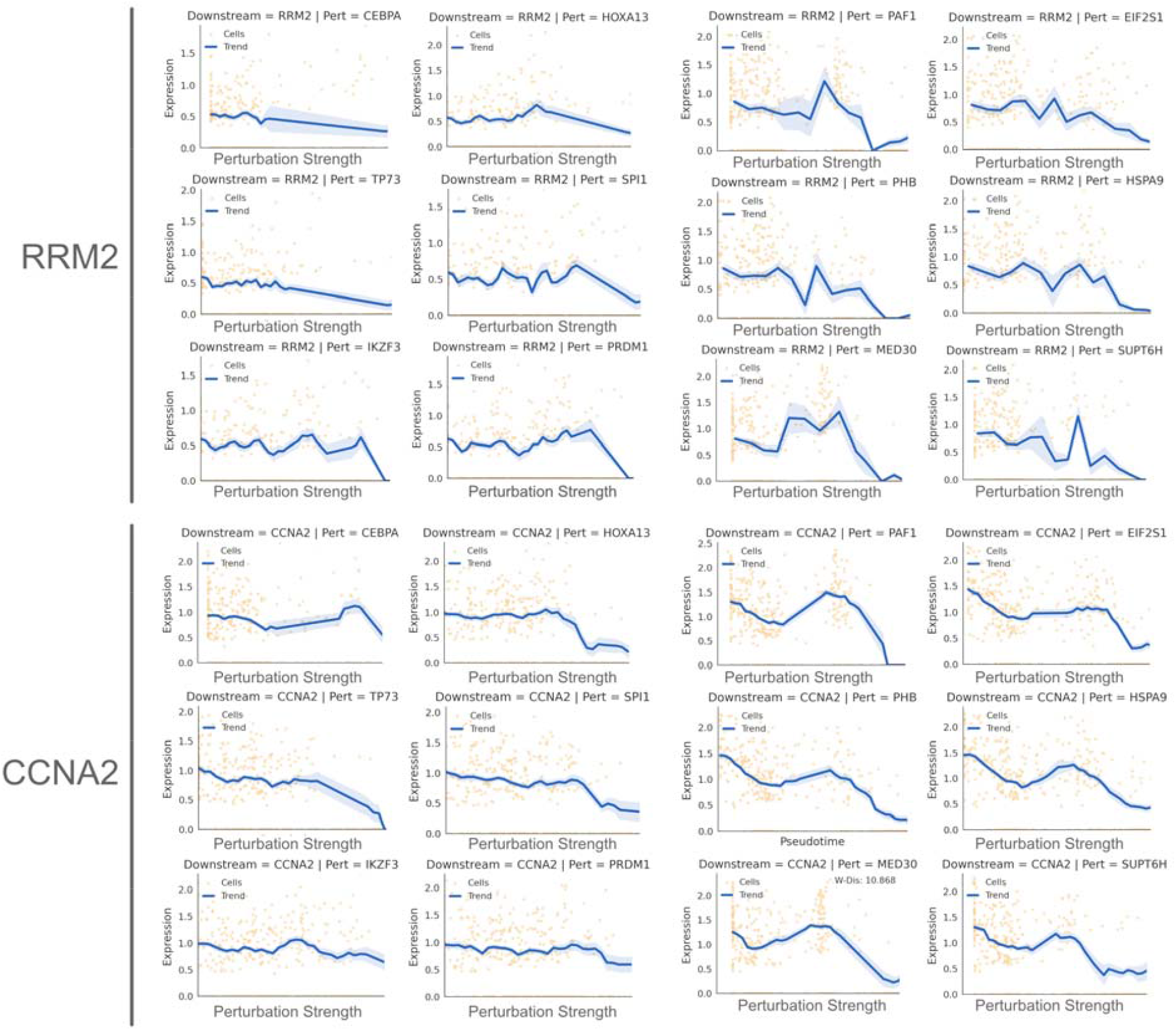
Universal downstream□gene response patterns across CRISPRa and CRISPRi. Threshold-type response of cell-proliferation genes *RRM2* and *MKI67*.

**Supplementary Figure 7.**
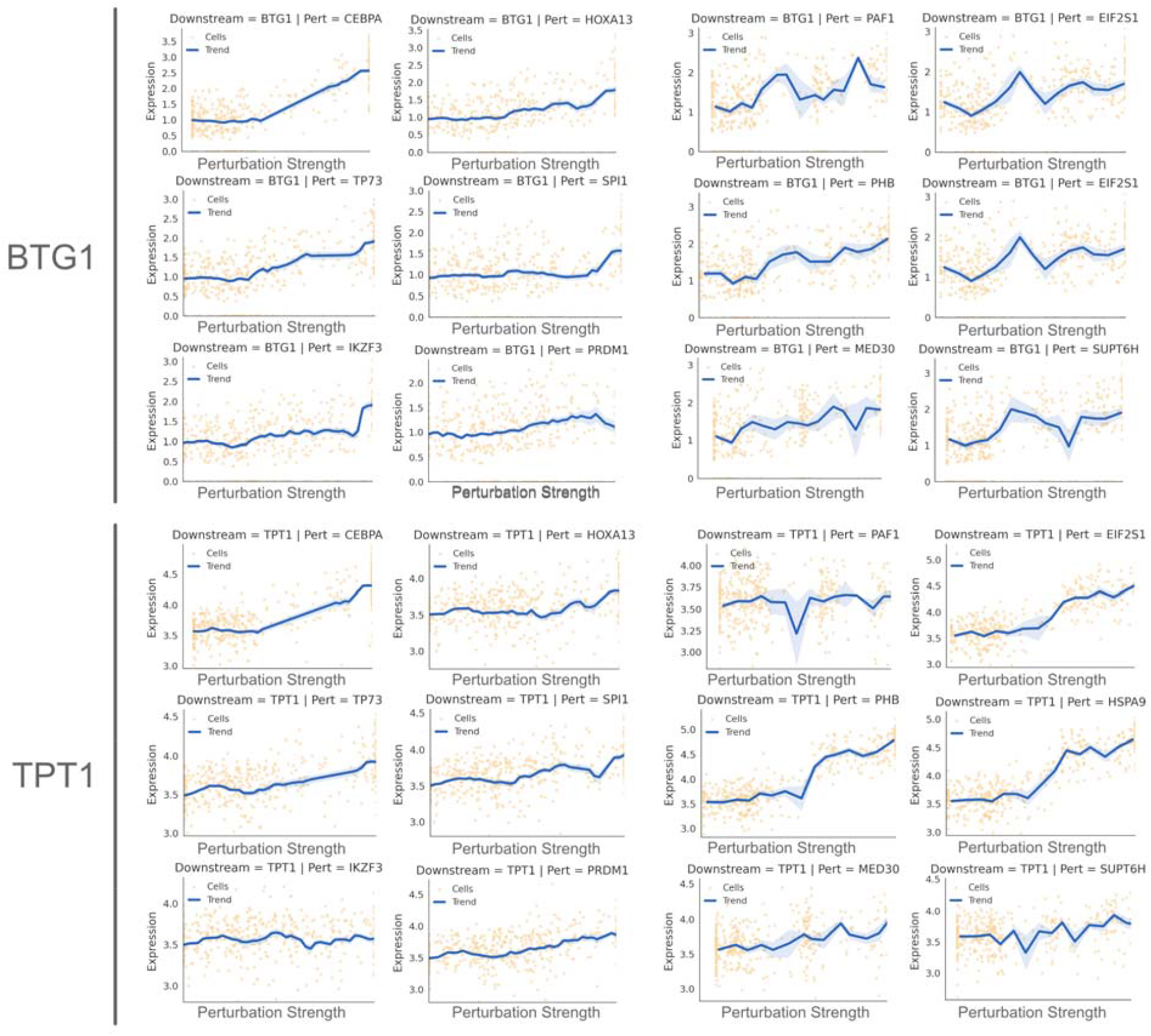
Universal downstream□gene response patterns across CRISPRa and CRISPRi. Proportional-type up-regulation of anti-apoptotic genes *BTG1* and *TPT1*.

**Supplementary Figure 8.**
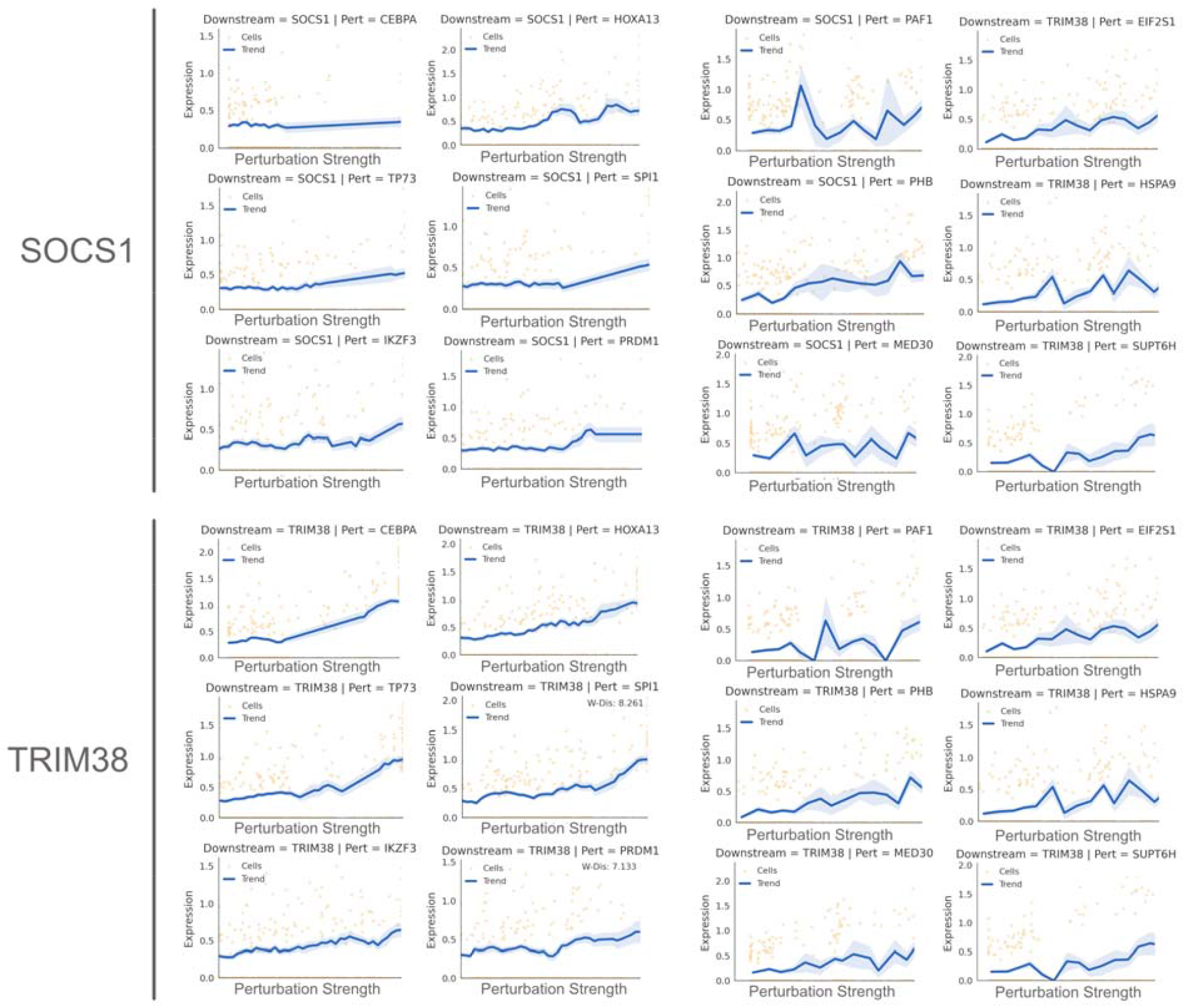
Universal downstream□gene response patterns across CRISPRa and CRISPRi. Proportional-type up-regulation of viral-response and infection-mechanism genes *SOCS1* and *TRIM38*.

**Supplementary Figure 9.**
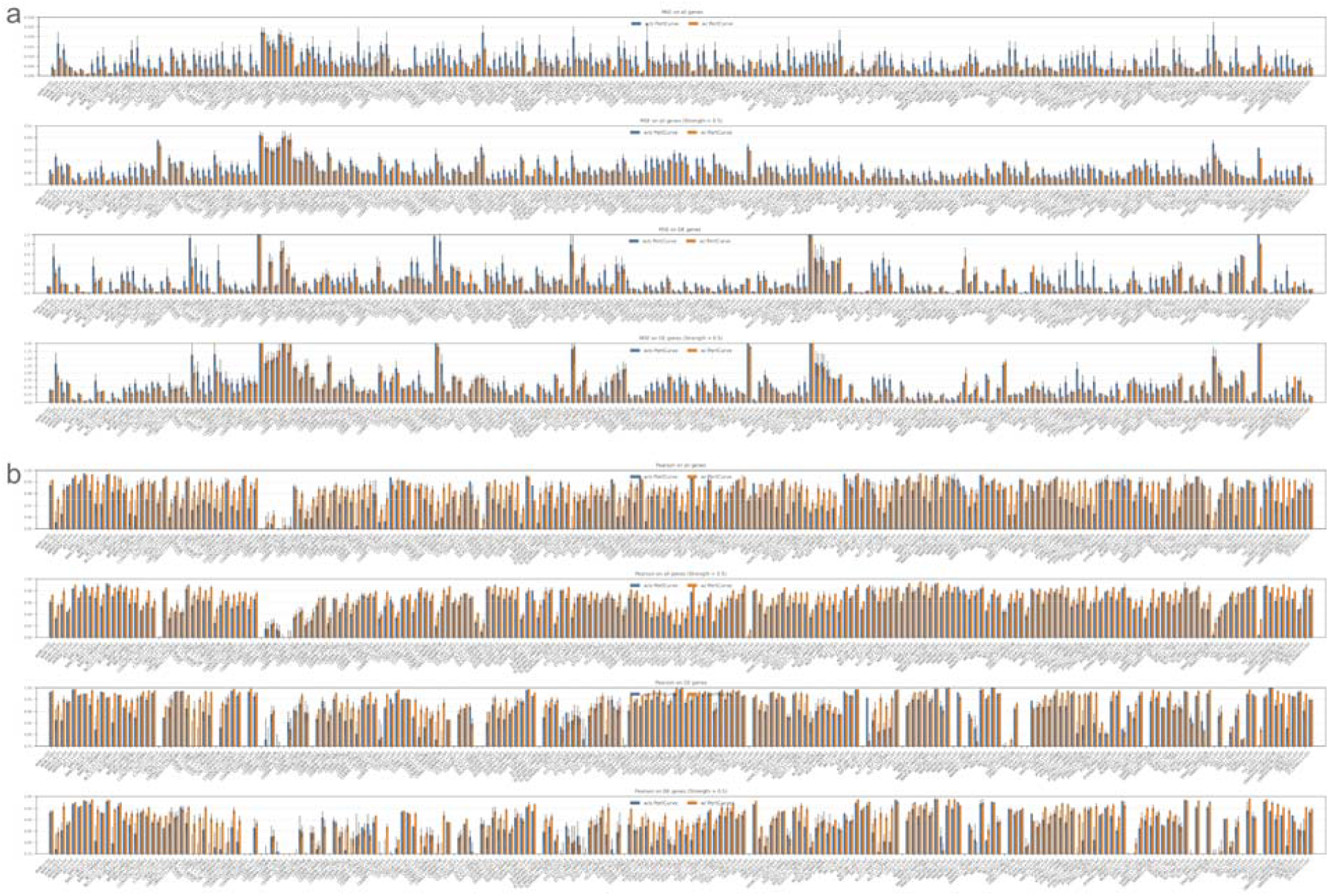
Performance evaluation and robustness metrics for GEARS on the Norman dataset in leakage-free evaluation settings. (a). Mean squared error (MSE) analysis across independent runs. Detailed MSE evaluation for individual perturbations across 10 independent runs of GEARS, assessed strictly on the holdout test split. Performance is stratified by overall gene expression (All genes MSE) and differentially expressed genes (DE genes MSE), evaluated across both the full cell population and the subset exhibiting phenotypic changes. Error bars denote standard deviation (s.d.) across the 10 replicates. (b). Pearson correlation metrics. Comprehensive assessment of model performance across all perturbations quantified by the Pearson correlation coefficient, under identical benchmarking conditions as in (a).

**Supplementary Figure 10.**
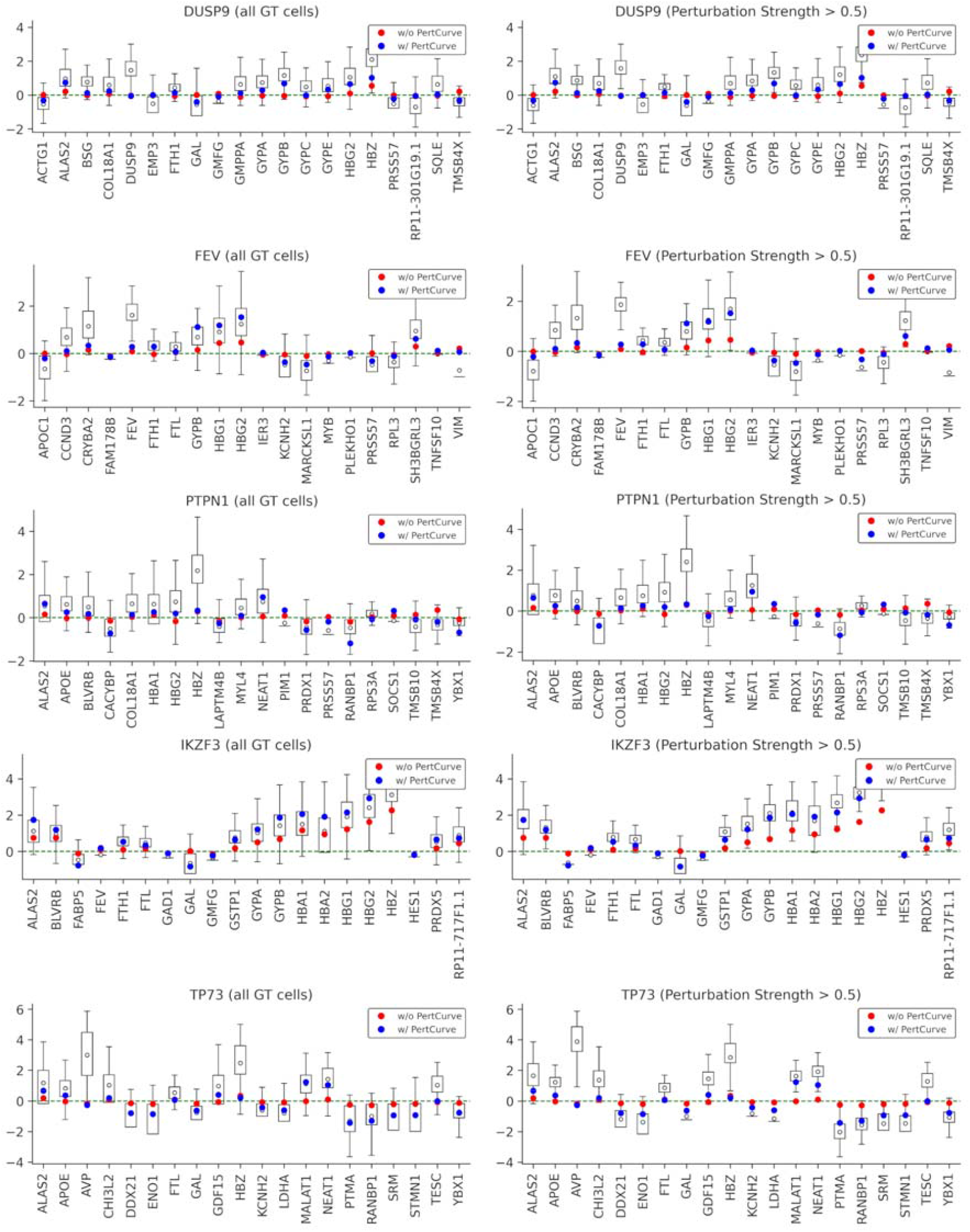
More examples of PertCurve enhancing predictive performance and biological fidelity in transcriptional response modeling. Gene-level prediction fidelity for *DUSP9, FEV, HOXA13, IKZF3* and *KLF1* perturbations. Boxplots show ground-truth single-cell expression distributions for downstream top20 DE genes (alphabetically sorted), with deterministic model predictions overlaid for the original GEARS model (*red*) and the PertCurve-enhanced model (*blue*).

